# Svep1 orchestrates distal airway patterning and alveolar differentiation in murine lung development

**DOI:** 10.1101/2021.07.26.453586

**Authors:** N Foxworth, J Wells, S Ocaña-Lopez, S Muller, P Bhayani, J Denegre, K Palmer, W Memishian, T McGee, SA Murray, PK Donahoe, CJ Bult, M Loscertales

**Affiliations:** Pediatric Surgical Research Laboratories, Massachusetts General Hospital, Boston, MA 02114; The Jackson Laboratory, Bar Harbor, ME 04609; Department of Surgery, Harvard Medical School, Boston, MA 02114

## Abstract

Disruptions in airway branching or alveolar differentiation during lung development can lead to severe respiratory deficiencies and neonatal death. The molecular mechanisms governing branching patterning and early alveolar formation remain elusive. Loss of *Svep1* function in mice results in various developmental defects, including lung hypoplasia and perinatal lethality. Our examination of the lungs of *Svep1* knockout (*Svep1^-/-^)* mouse embryos, both *in vivo* and *in vitro*, revealed that *Svep1* mutants exhibit an increase in the number of disorganized distal airway tips and progressively greater disruption of lung lobe morphology over time and saccular development. *Svep1* interacts with FGF signaling to regulate smooth muscle differentiation and, together with *Fgf9,* guides airway branching patterning. Transcriptomic data from the lungs of *Svep1^-/-^* embryos revealed dysregulated gene expression affecting saccular maturation. Our findings demonstrate that *Svep1* is a key extracellular matrix player shaping airway morphology and influencing alveolar fate. These insights offer potential avenues for therapeutic interventions in congenital lung disorders.

## Introduction

The architectural blueprint of the lungs emerges from two fundamental developmental processes: branching morphogenesis, responsible for constructing the intricate airway tree structure, and differentiation, which generates specialized cells in distinct lung regions, predominantly after the formation of the tracheobronchial tree ^1^. The cessation of molecular genetic signaling promoting branching is a prerequisite for distal alveolar cell differentiation ^2, 3^.

In mice, lung development begins at embryonic day 9.5 (E9.5) ^4, 5^ when two primary buds emerge from the ventral foregut. During the pseudoglandular stage (E12.5 – E16.5), epithelial progenitors at the bud tips undergo an iterative process of budding and bifurcation into the surrounding mesenchyme and this is followed by expansion of the diameter and length of the respiratory tree during the canalicular stage (E16.5-E17.5). During the saccular stage (E17.5-P5), the vasculature develops along the airway and terminal airway tips widen. The alveolar epithelium subsequently differentiates into alveolar type 2 (AT2) cells, responsible for surfactant protein C (SFPC) production, and the thin, gas-exchanging alveolar type 1 (AT1) cells. Primitive septa arise at this stage. Secondary septation and alveolar maturation occur during the postnatal alveolar stage (P5-P30) ^5^.

The geometry of the airway tree is generated by three local branching modes: domain, planar, and orthogonal ^6^. During domain branching, new branches grow perpendicularly around the circumference of the parent branch and generate the initial scaffold of each lobe and the overall shape of the lung. Planar bifurcation occurs at the tips of the airway epithelium and generates branches in the same plane, resulting in the thin edges of the lung lobes. Orthogonal bifurcation generates branches perpendicular to prior branches, thus filling in the interior of the lung. Trifurcations are rare and appear principally at the lung surface ^7, 8^. As branching proceeds, proximal-distal patterning is established, and cellular specification follows ^3, 9, 10, 11^. Lung bud tip progenitors persist and proliferate and subsequently undergo differentiation into alveolar type 1 (AT1) and alveolar type 2 (AT2) cells ^3^. The cells of the primal airway stalk differentiate into multiple proximal cell types ^12, 13, 14, 15^.

Studies in laboratory mice have shown that branching morphogenesis requires mesenchymal-epithelial interactions, and various signaling pathways, particularly the FGF pathway, play important roles in branching ^16^. Fibroblast growth factor 10 (FGF10) is secreted by mesenchymal cells and acts via the *FGFR2* receptor to maintain a permissive environment for branching morphogenesis. FGF10 maintains distal epithelial SOX9^+^ progenitor cells in an undifferentiated state. These cells play a pivotal role in subsequent alveolar and mesenchymal differentiation ^17, 18, 19, 20, 21, 22, 23, 24, 25, 26, 27, 28^. FGF10^+^ cells give rise to bronchial airway smooth muscle cells during early branching ^29, 30, 31, 32^. FGF9 is also required for lung development and plays different roles depending on its spatial and temporal expression. In the mesothelium, FGF9 plays a primary role in mesenchymal growth; in the epithelium, FGF9 controls epithelial branching ^33, 34, 35, 36^. The extracellular matrix (ECM) also plays important roles in epithelial branching and alveolarization ^37, 38, 39, 40, 41, 42^ and contributes to organ shape ^43^.

*Svep1* (Sushi, Von Willebrand Factor Type A, EGF and pentratix domain-containing 1), also known as Polydom, encodes an ECM protein that is ubiquitously expressed in the tissues of adult mice and plays roles in cell adhesion ^44^, epidermal differentiation ^45^, and lymphatic vessel formation and remodeling ^46, 47^. *Svep1* homozygous mutant mice die at birth with edema and body abnormalities ^45, 47, 48, 49^. We investigated the role of *Svep1* in the development of lungs of mice homozygous for the *Svep1^tm1b(EUCOMM)Hmg^* allele (hereafter, *Svep1^-/-^*) generated by the Knockout Mouse Phenotyping Program (KOMP2) program ^49^. We observed that the loss of *Svep1* disrupts the basic lung branching program resulting in irregular lung lobe shape due to an aberrant increase in distal airway tips. Smooth muscle differentiation is also disrupted in the lungs of *Svep1^-/-^* embryos. Through the treatment of lung explant cultures with exogenous SVEP1 protein, we demonstrated that *Svep1* functions as an inhibitor of branching morphogenesis. Furthermore, we demonstrated that inactivation of FGF signaling recovers smooth muscle differentiation in *Svep1^-/-^* lungs and that exogeneous FGF9 treatment restores SVEP1 branching inhibition, implying a potential role for SVEP1-FGF9 signaling specific to branching.

## Results

### Svep1 is expressed in lung mesenchyme with high expression at the lung periphery

*Svep1* is expressed in the lung mesenchyme, especially at the tip and edges of the lung (Fig. 1a-*d* and Suppl. Fig. 1a-b). *Svep*1 mRNA expression was strong in the mesenchyme adjacent to the distal epithelial airways and around bifurcating tips (Fig.1a and Suppl. Fig.1a). We detected SVEP1 protein expression on the epithelium membrane (ECAD+) of very distal lung buds (Fig.1b and Suppl. Fig. 1b) and microvasculature (Suppl. Fig.1c). At the saccular stage (E18.5), transcript and protein expression of *Svep1* was detected in the primary septa of lung parenchyma and in the mesenchyme next to the proximal epithelium (Fig.1d and Suppl. Fig.1d). Consistent with previous findings, *Svep1* expression during lung development was high between E14.5 and E18.5 ^50^ (Suppl. Fig.1e). Single cell RNA-seq data demonstrated that expression of *Svep1* in the lungs of E18.5 wild type embryos was prominent in matrix fibroblasts (Suppl. Fig. 3a).

**Figure 1.**
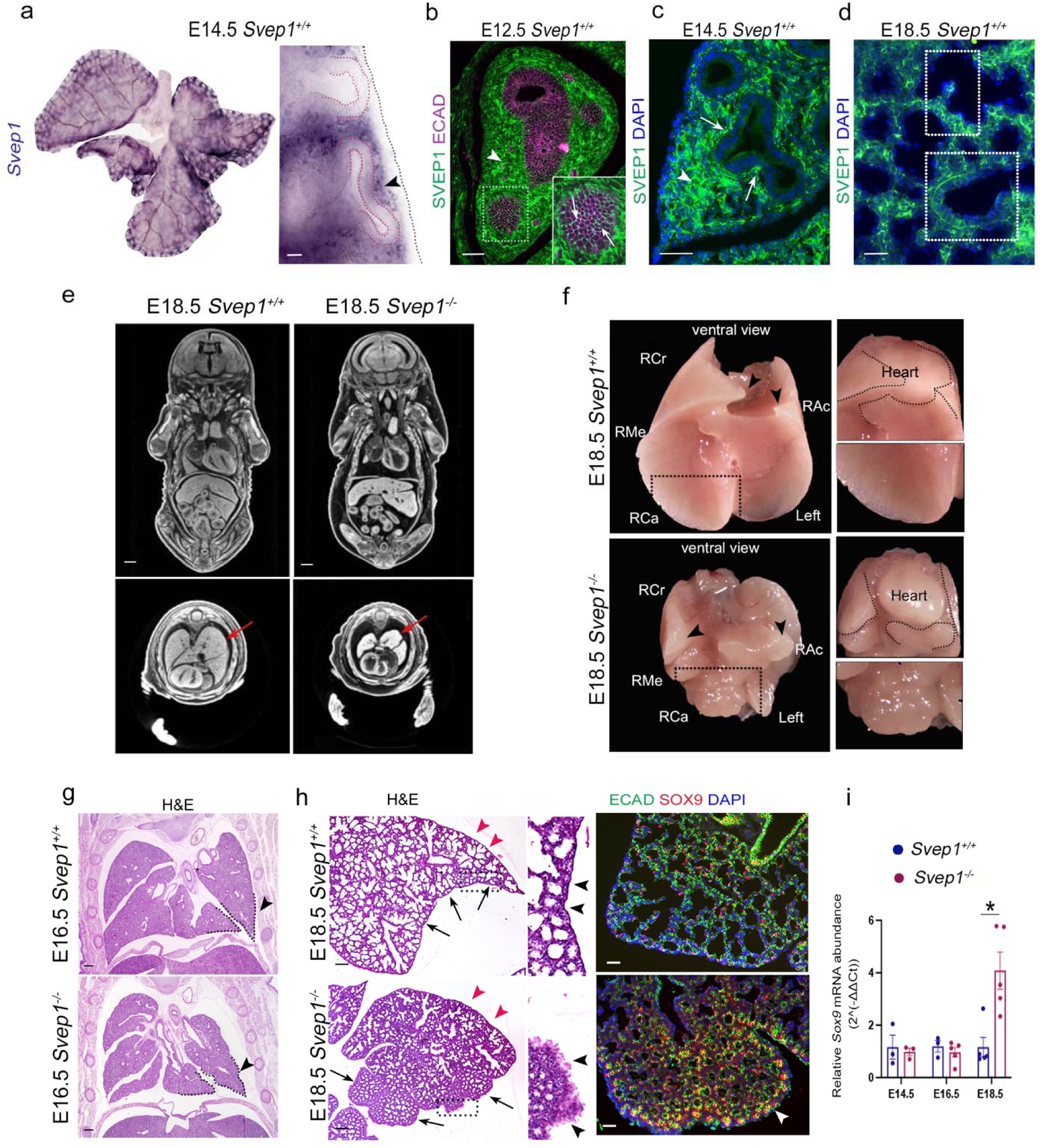
Svep1 expression pattern and Svep1-/- hypoplastic lung lobes anomalies. **(a)** *In situ* hybridization of lungs from E14.5 wild type embryos showed strong *Svep1* mRNA expression in the mesenchyme at the lung periphery and around bifurcating distal airway tips (right inset outlined in red, arrowhead). **(b)** Arrowhead shows Mesenchymal expression of SVEP1 protein in normal E12.5 lung. White arrows indicate the location of the SVEP1 protein on the epithelial membrane stained with ECAD (purple) in a distal airway bud. **(c)** SVEP1 expression is strong in the mesenchyme towards the lobe tip of the E14.5 lungs and branching cleft sites (arrowheads). **(d)** SVEP1 protein localizes in the lung parenchyma including the primary septa (upper box) and adjacent to the proximal airway epithelium (lower box). **(e)** microCT scan of E18.5 embryos showing multiple defects including thinning of the diaphragm and lung hypoplasia (red arrows). **(f)** Ventral view of E18.5 of *Svep1^+/+^* and *Svep1^-/-^* lungs. The arrowheads indicate the tips of the lung lobes, which are short and rounded (highlight in black) in *Svep1^-/-^* mice and have like a cauliflower morphology. **(g)** H&E-stained lung sections embryos demonstrate that E16.5 lungs from *Svep1^-/-^* embryos are smaller and have irregular lobe edges. Black arrowheads indicate left lobes with highlighted surface. **(h)** The red arrowheads show the dorsal side of the E18 lungs, whereas the black arrows highlight the ventral side. *Svep1^-/-^* embryos show defective saccular development mainly at the edge on the ventral side of the lung (inserts arrowheads). Co-localization of ECAD and SOX9 confirms high expression of SOX9 in distal airways (white arrowheads) in the lung of *Svep1^-/-^* embryos at E18.5. **(i)** Plots showing *Sox9* relative mRNA expression at E14.5, E16.5, and E18.5. mRNA expression is significantly high at E18.5 in *Svep1^-/-^* embryos (average number of tips per area ± SEM; p < 0.05; n ≥ 5). Right cranial (RCr), Right Caudal (RCa), Right Medial (RMe), and Right Accessory (RAc) lobes. *Scale bars: 50μm (a, b, c, d), 100 μm*.

### Hypoplastic and aberrant lung shape in Svep1^-/-^ embryos

Consistent with previous reports ^49^, *Svep1^-/-^* embryos displayed perinatal lethality, whole-body edema, and lung hypoplasia that was prominent at E18.5 by microCT (Fig. 1e and Suppl. Fig. 1f). *In situ* hybridization confirmed the absence of *Svep1* expression in the lungs of homozygous knockout animals (Suppl. Fig. 1g). Examination of gross examination of embryos at E18.5 revealed small lungs with lobes that failed to enclose the heart and had a highly lobulated, cauliflower-like appearance (Fig. 1f). Histological analysis at E16.5 showed small lungs in mutants with irregular shape; at E18.5, small saccules were observed along a dorsal-ventral gradient, accompanied by a reduced lumen primarily on the ventral side (Fig. 1g,h). Noticeable differences, including a thickening along the lobulated ventral edges of the lung composed of mesothelium, were observed in *Svep1^-/-^* embryos compared to wild type embryos reflecting developmental abnormalities in the lung mesothelium of mutant mice (Fig. 1h).

### Saccular development defects in Svep1^-/-^ embryos

Sacculation begins in the murine lung shortly after E16.5, with the flattening of AT1 cells at the proximal-to-distal opening of the distal airway lumen. Cells in the developing distal airways at the lung edge are the last to differentiate ^11, 30, 51^. Examination of distal airway morphology of *Svep1^-/-^* embryos at E18.5, through labeling the lung epithelium with ECAD and the distal progenitor marker SRY-box transcription factor 9 (SOX9), revealed that the densely packed distal airways at the ventral edge exhibited a morphology typical of the earlier pseudoglandular stage instead of the saccular stage (Fig. 1h). Quantitative real-time PCR (qPCR) confirmed elevated expression of *Sox9* in the lungs of *Svep1* knockout embryos at E18.5 but not at earlier time points (Fig. 1i). Co-staining of SOX9 with the cell proliferation market KI67 showed proliferating epithelial progenitor cells in E18.5 lungs, mainly at the lung periphery in *Svep1^-/-^* mutants (Suppl. Fig. 1h). Our results indicate that the distal epithelium in *Svep1* lungs is in an earlier developmental stage, characterized by a higher proportion of epithelial cells in a progenitor stage, rather than in the saccular period when cells are differentiating into specialized lung epithelial type I and II cells. Our finding highlights the essential role of Svep1 in the maturation of the distal lung epithelium.

### Lungs of Svep1^-/-^ embryos show an aberrant increase of distal airways altering lung lobe shape

To determine whether *Svep1^-/-^* embryos have branching defects, we stained E16.5 lungs with SOX9 and ECAD which revealed densely packed distal airways in *Svep1^-/-^* embryos. Mutant lobe tips had multiple distal branches and rosettes on the ventral lobe side (Fig. 2a). To further examine defects in the branching program modes^6^ (Fig.2b) in *Svep1* mutant lungs, we stained whole lung and individual lobes with SOX9 at E13.5, E14.5, and E15.5 and showed that *Svep1* branching defects accumulate over time (Fig. 2c-e and Suppl. Fig. 2a,b). We observed branching and lobe shape anomalies in *Svep1^-/-^* lungs as early as E13.5, primarily at the lung tips and edges (Fig. 2c,d). Moreover, the branching abnormalities observed in the mutants were not uniform across the lung lobes, leading to different outcomes in the shape of the lobes. At E13.5 for example, the shape of the Right Caudal (RCa) lobe was narrower in *Svep1^-/-^* embryos compared to wild-type embryos. The Right Accessory (RAc) and Right Medial (RMe) lobes of *Svep1* mutant embryos terminated in two or more buds instead of a single tip typically observed in lungs of wild type embryos (Fig. 2c). Further examination of the left lobes showed that *Svep1^-/-^* embryos at E13.5 exhibited peripheral branch mis-localization and additional budding, disrupting the normal planar bifurcation of the lung (Fig. 2d and Suppl. Fig. 2a). When planar bifurcation is disrupted at the lobe tip, ectopic branching angles occur at the lung tips in mutants (Fig. 2d). This observation suggests that *Svep1* deficiency impairs lung branching angles early in development.

**Figure 2.**
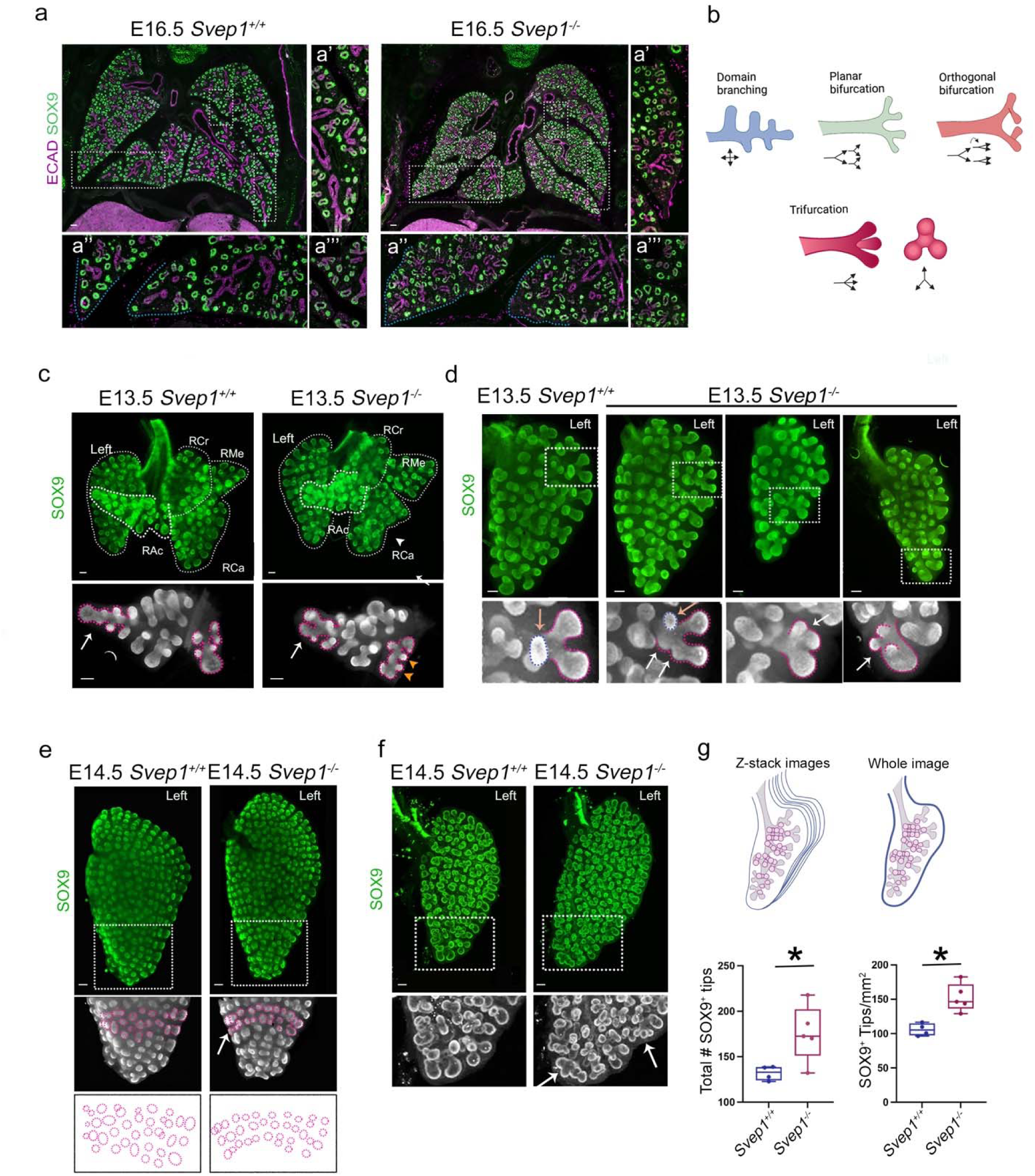
Airway organization defects and abnormal branching patterns in *Svep1^-/-^* embryonic lungs resulting in lung lobe shape anomalies. **(a)** Co-localization of epithelial (ECAD) and epithelial progenitor (SOX9) markers showing distal airways in lungs of E16.5 mice. **(a’)** The blue highlights in inserts represent the distal tip of the normal and mutant RAc lung lobes, with the mutant left lobe showing multiple branching. **(a’’)** Inserts illustrate airway density at the lung periphery in WT and mutants, with mutants having higher density. **(a’’’)** In *Svep1^-/-^* lungs, distal airway tips are organized in packed airway rosettes on the ventral side of the lung lobe. **(b)** Illustration of the three branching modes that pattern the airways of the lung (domain branching, planar bifurcation, orthogonal bifurcation) and distal trifurcation. **(c-f)** Representative Z-stack images show SOX9 immunofluorescence **(c)** E13.5 Whole lungs of normal and *Svep1^-/-^* embryos. The images show *Svep1^-/-^* lungs with abnormal RCa lobe morphology (arrowheads), RAc lobe distal tip branching (arrows), and secondary branching pattering (orange arrowheads). **(d)** Pictures of E13.5 *Svep1^+/+^* and *Svep1^-/-^* embryos showing ectopic branching/budding anomalies in mutant embryos (areas outlined in pink in lower inserts). **(e)** E14.5 left lung lobes of *Svep1^+/+^* and *Svep1^-/-^* embryos. SOX9^+^ buds highlighted in pink and cartoon show airway tip arrangement in normal and mutants. Arrow shows defective tip arrangement resulting in abnormal lung tip morphology in *Svep1^-/-^* embryos. **(f)** Cleared tissue immunofluorescence of SOX9 in the left lung lobe at E14.5 reveals airway disorganization in *Svep1^-/-^* embryos vs *Svep1^+/+^* embryos. Distal lung tips highlighted show branching trifurcations in the *Svep1^-/-^* embryo (white arrow) and aberrant distal tip airway. **(g)** Representation of the left lung lobe Z-stack for distal airway quantification. Whole lungs of *Svep1^-/-^* embryos at E14.5 have a statistically significant increase in the total number of SOX9 positive airway tips and ratio of tip and area compared to wild type embryos (average number of tips for area ± SEM; p < 0.05; n ≥ 5). *Right cranial (RCr), Right Caudal (RCa), Right Medial (RMe), and Right Accessory (RAc) lobes. Scale bars: 100μm (a-e)*.

The orthogonal bifurcation mode of the branching program fills the lung’s exterior and interior surfaces in a specific pattern where each new bud maintains its boundaries with others and creates the surface of the lung lobes ^6^. However, this orthogonal pattern was disrupted in mutant embryos at E14.5, with the airways rearranged into a “row of beads” pattern, resulting in abnormal rounded tips (Fig. 2e). Moreover, the left lung lobes of E14.5 mutant embryos were 8% more elongated than those of wild type *Svep1* embryos (length/width ratio (mm) *Svep1^+/+^* =1.60; *Svep1^-/-^* =1.90 *p <0.001 t test;* Suppl. Fig. 2c). Tissue cleared left lobes stained with SOX9 at E14.5 confirmed abnormal mutant left lobe morphology as well as multi budding of the distal airways and peripheral trifurcations (Fig. 2f).

At E15.5, aberrations persist in lung branching patterns along the margins and ventral side, leading to substantial alterations in lobe elongation in *Svep1^-/-^* embryos (Suppl. Fig. 2b,c,d). Staining left lung lobes of *Svep1^-/-^* embryos at E15.5 with DAPI (4’,6-diamidino-2-phenylindole) revealed prominent abnormalities on the lung surface (Suppl. Fig. 2e).

Quantification of SOX9+ in a whole lung Z-stack imaging revealed a significant increase in both the total number of SOX9+ distal tips and the ratio of tips to area by E14.5 (Fig. 2g; p < 0.05 t-test). Overall, these findings demonstrate that *Svep1* is crucial for normal branching morphogenesis, and its absence leads to disorganization of airway patterning, primarily affecting the tips, edges, and ventral areas of the lung, which in turn alters the overall shape of the lung lobes.

### Lungs of Svep1^-/-^ embryos have ectopic branching, decreased planar bifurcations, and increased trifurcations

To investigate which of the main branching programs are affected in *Svep1* mutants, we modeled the branching patterns of normal and mutant lungs. We labeled tissue-cleared whole lungs from E14.5 embryos with ECAD and SOX9 and generated a 3D reconstruction composed of Z-stack images. We observed significant structural differences in the airways of wild type and *Svep1^-/-^* lungs, confirming the disorder of the developing airway tree as well as the aberrant duplication of the distal airway tips in *Svep1^-/-^* embryos (Fig. 3a).

**Figure 3.**
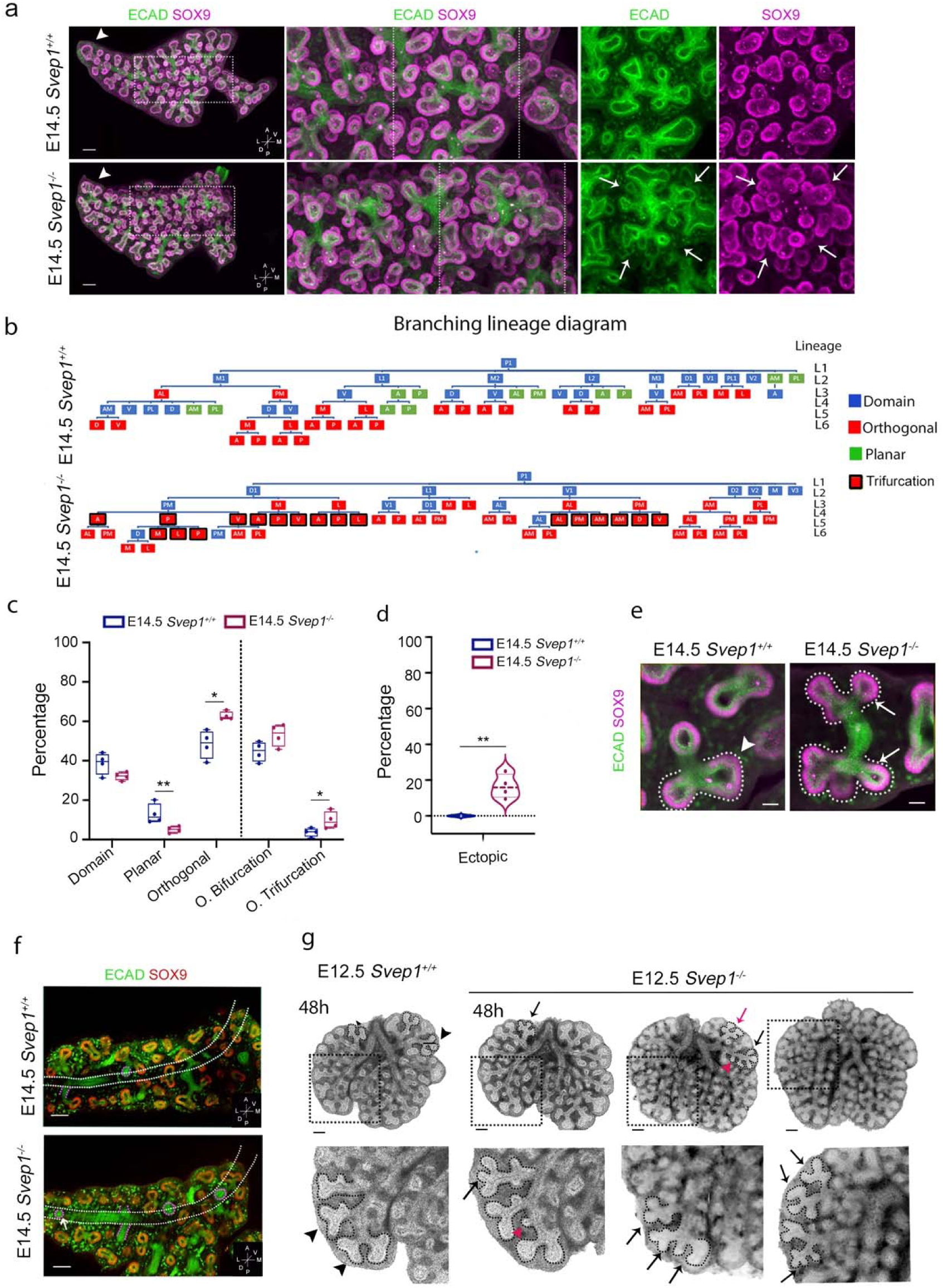
Disrupted branching programs and altered airway architecture in *Svep1^-/-^* RAc lung lobes at E14.5. **(a)** Compilation of a Z-stack image series for tissue-cleared whole RAc lung lobes labeled with ECAD (green) and SOX9 (purple) highlight disorganized branching in *Svep1^-/-^* embryos at E14.5. Arrows (lower right) point to branching airways with rosette-like arrangements and ectopic airway tips growing in random directions. Arrowheads (left panels) at the RCa lobe tips indicate a single tip in the lobe of *Svep1^+/+^* embryos but multi branched tips in *Svep1* knockout embryos. **(b)** Schematic illustrating representative domain (blue), planar (green), and orthogonal (red) branching program types of *Svep1^+/+^* and *Svep1^-/-^* RAc lobes. Lungs of *Svep1^-/-^* embryos lack planar bifurcations and have many trifurcating branch tips (red highlighted in black). **(c-d)** Graphs displaying the percentages of branching modes, orthogonal bifurcations and trifurcations **(c)** and ectopic budding **(d)** in E14.5 Rac lobes of wild type and *Svep1^-/-^* embryos. **(e)** Representative single Z-stack images of RAc lobes at E14.5, stained with ECAD (green) and SOX9 (purple), show bifurcations (white dotted lines and white arrowhead) in *Svep1^+/+^* embryos, but trifurcations (white dotted lines and white arrows) in *Svep1^-/-^* mice. **(f)** Single Z-stack image of RAc lobes stained with ECAD (green) and SOX9 (red) display ectopic domain branching in *Svep1^-/-^* mice (arrow). **(g)** Lung explants from E12.5 embryos after 48 hours in organ culture. Explants from *Svep1^+/+^* embryos show normal bifurcation (black arrowheads) whereas explants from *Svep1^-/-^* embryos show trifurcations (black arrows) and ectopic budding (red arrowheads). (Average number of tips per area ± SEM; p < 0.05; n ≥ 5) (c,d). *Scale bars 100μm (a, g); 50μm (e)*.

To quantify the differences in branching morphology in the lungs of mutant vs wild type embryos, we generated a branching lineage diagram for the right accessory (RAc) lobe at E14.5. Each branch was characterized as one of three branching modes: domain, planar, or orthogonal ^6^. Branches and direction of growth were mapped to create a lineage diagram in which trifurcations were also indicated (Fig. 3b). Orthogonal branching was significantly increased in mutants, contributing 63% of all branches compared to 49% (p < 0.01, t-test) in the lungs of *Svep1^+/+^* embryos (Fig. 3c). In contrast, planar branching in the RAc lobes was significantly decreased in mutant (5%) compared to wild type (13%) embryos (p < 0.05, t-test; Fig. 3c). Trifurcations were more prevalent in *Svep1^-/-^* lungs (10% of all branches) compared to the lungs of *Svep1^+/+^* embryos (4% of all branches) (p < 0.05 t-test; Fig. 3c and Fig.3e). An average of 17*%* of first-level domain branches originating from the primary bronchus were ectopic in *Svep1*^-/-^ lungs, while no ectopic branching was observed in the lungs of wild type embryos (Fig. 3d and Fig. 3f). In RAc lobes of wild-type lungs, daughter branches of the primary bronchus sprouted from one of only three directions: anterior, ventral-posterior, and dorsal-posterior. In all the *Svep1^-/-^* lungs analyzed, daughter branches sprouted from the primary bronchus in random directions (Fig. 3f). In summary, loss of *Svep1* disrupts airway patterning through decreased planar bifurcations, increased trifurcations, and ectopic domain branching that arise in random directions.

To determine if the ectopic and trifurcated branches in *Svep1^-/-^*lungs was the result of loss of *Svep1*, we analyzed time-lapse images of E12.5 lung explants for 48 hrs. (Fig. 3g, Video1, Video2, Video 3). We observed ectopic branching arising from the main and secondary branches and trifurcations at the lung tips. Taken together, these results suggest that *Svep1* controls the position and orientation of new airway epithelial buds, defines distal bifurcations regulating the form and structure of the growing epithelium, and is, therefore, critical for determining the bronchial tree architecture and, ultimately, the final shape of the lobes of the lung.

### SVEP1 recombinant protein inhibits airway branching in normal E12.5 lung explants and restores normal branching in Svep1^-/-^ lungs

The increased branching observed in the distal airways of *Svep1^-/-^* embryos suggests that SVEP1 normally acts as an inhibitor of lung branching morphogenesis and is essential for regulating lung airway budding. To further investigate the function of *Svep1* in branching morphogenesis, we treated lung explants of wild type E11.5-E12.5 embryos for 48hrs with a recombinant SVEP1 (rSVEP1) protein. Time-lapse images revealed significantly decreased branching in the treated lung explants (Fig. 4a; Video 4 and Video 5), quantified at the peripheral airway tips (Fig. 4b). SVEP1 branching inhibition was more profound at earlier stages (11.5) (data not shown). When lung explants from E12.5 *Svep1^-/-^* embryos were treated with rSVEP, distal ectopic and trifurcated branches were reduced, and observed in only one of six explants (Fig.4c). The *other Svep1^-/-^* explants exhibited distal bifurcation that resembled wild type lungs (Fig.4c).,

**Figure 4.**
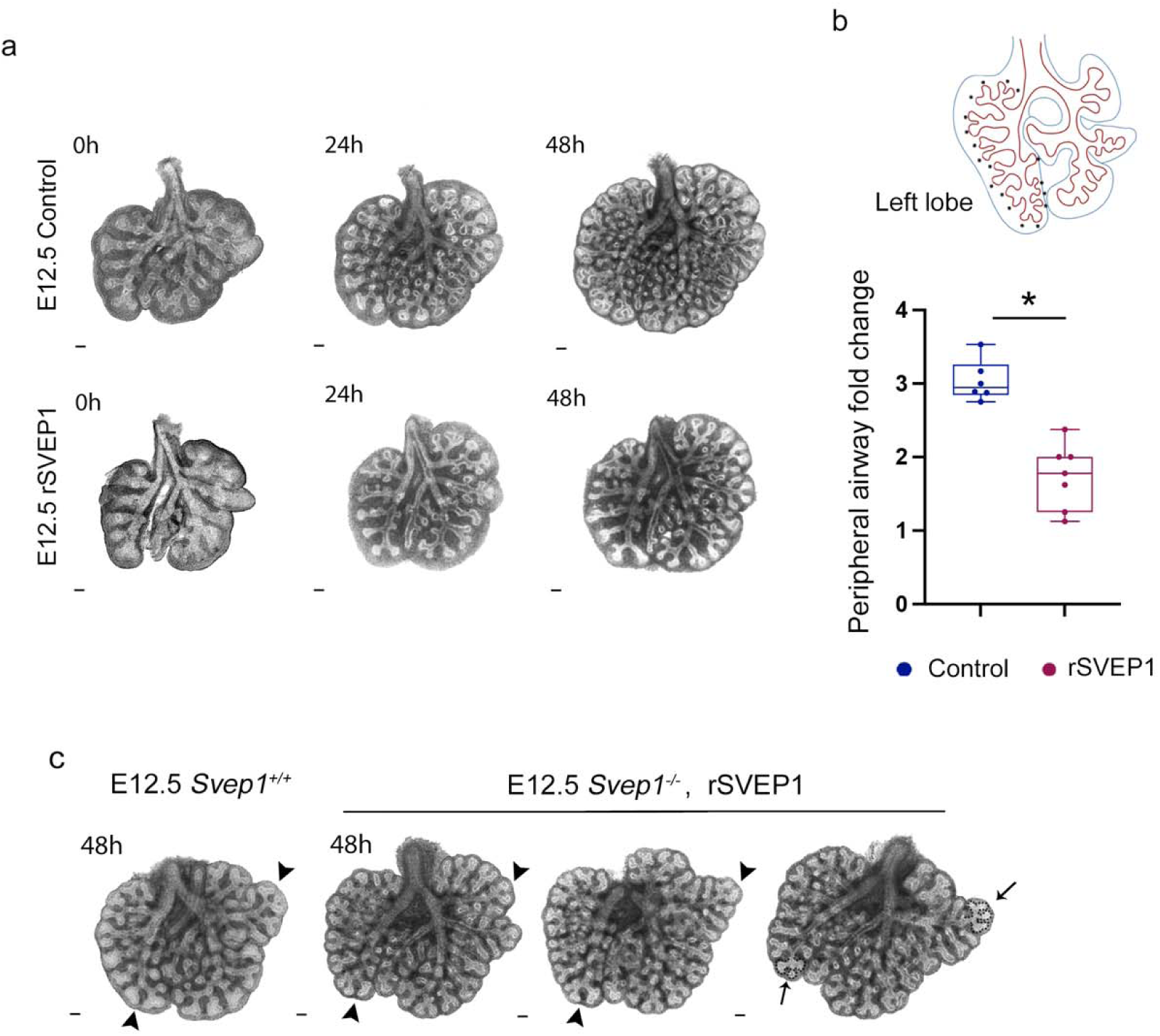
SVEP1 treatment modulates airway branching in E12.5 lung explants. **(a)** Images of lung explants from E12.5 *Svep1^+/+^* embryos cultured for 48 hrs. Images of lung explants from E12.5 *Svep1^+/+^* embryos cultured for 48 hrs. show a significant reduction of branching (control, top row) when treated with SVEP1 peptide (bottom row). **(b)** Plot showing peripheral airway counts of a left lung lobe after 48 hrs cultured with and without SVEP1 treatment. **(c)** *Svep1^-/-^* lung explants treated with SVEP1 for 48 hours show a reduction in trifurcations compared to untreated lung explants. Black arrowheads indicate bifurcation; arrows indicate trifurcations. (Average counts ± SEM; *t* test p < 0.05 n≤ 6). *Scale bars 100 µm* (a,c).

### The loss of Svep1 leads to defects in smooth muscle and bronchial airway development

To characterize further the lungs of *Svep1^-/-^* embryos at the saccular stage, we conducted RNA sequencing (RNA-seq) of whole lungs of *Svep1^+/+^* and *Svep1^-/-^* embryos. Term enrichment analysis of the functional annotations for the differentially expressed genes revealed that genes associated with cell mobility and adhesion, epithelial proliferation, extracellular matrix organization, and collagen binding were significantly downregulated in *Svep1^-/-^* lungs (Fig. 5a and Fig. 5b).

**Figure 5.**
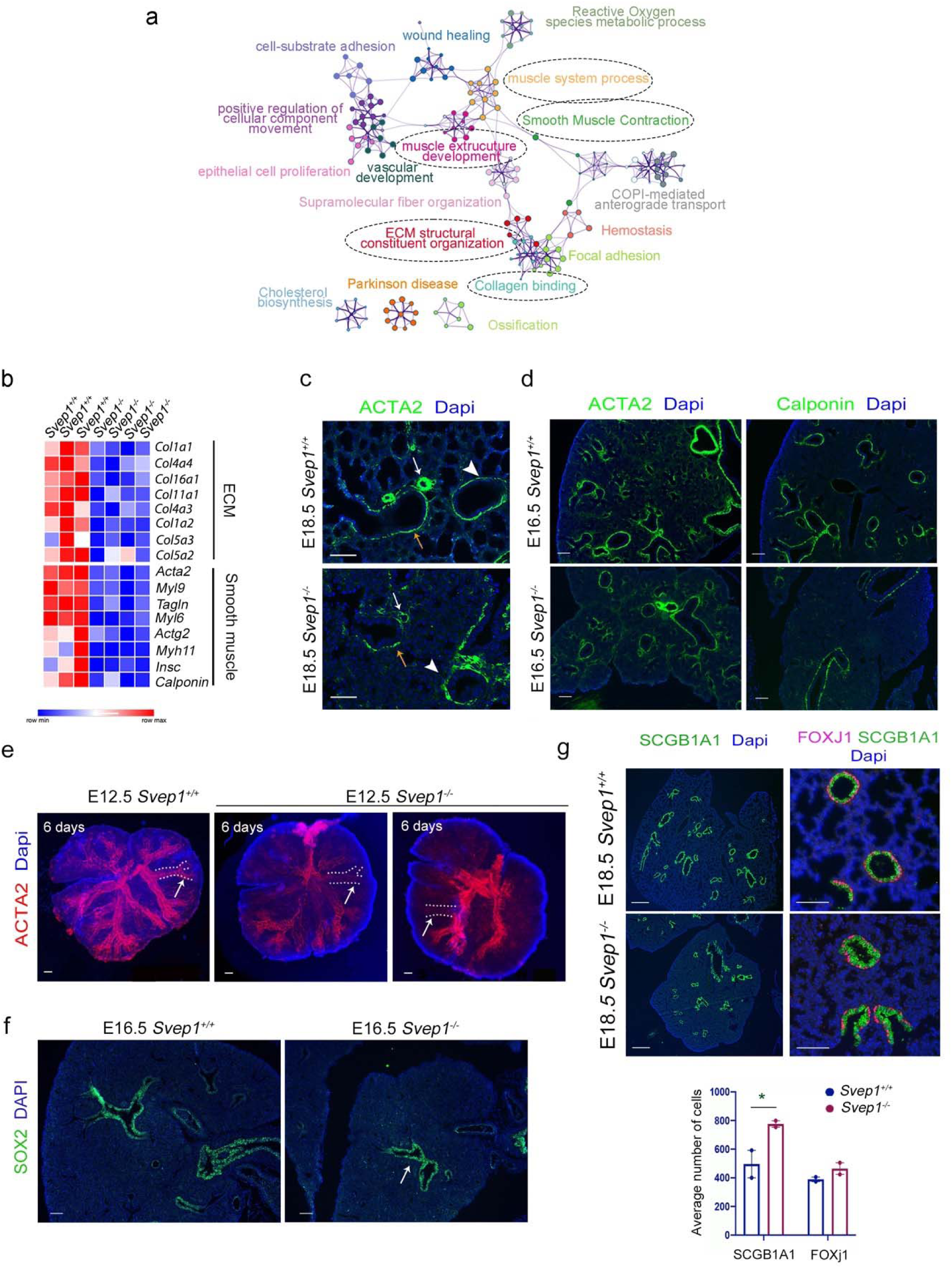
Impaired smooth muscle and airway epithelium differentiation in *Svep1^-/-^* embryonic lungs. **(a)** Interactome of downregulated genes in lungs of E18.5 *Svep1^-/-^* embryos with clusters labeled by enriched annotation terms. **(b)** Gene expression heatmap from RNA sequencing shows significantly downregulated genes in lungs from *Svep1^-/-^* embryos are associated with collagen and smooth muscle genes. **(c)** ACTA2 and Calponin staining at E16.5. **(d)** ACTA2 and calponin localization in lung at E18.5 (white arrowheads indicate distal respiratory bronchioles; orange arrows indicate proximal bronchioles; white arrows indicate blood vessels). **(e)** ACTA2 staining (red) in wild type and *Svep1^-/-^* E12.5 lung explants each cultured for six days, showed a reduction of smooth muscle cell formation in secondary and tertiary bronchi of the mutants (white arrows). **(f)** SOX2 Immunofluorescence staining marking the progenitor cells of the epithelium in the proximal airways. Arrow marks the proximal airway at the core of the *Svep1^-/-^* lung. **(g)** Immunofluorescence of the club cell marker SCGB1A1 (green) highlights the small proximal airways in mutant lungs. SCGB1A1 (green) colocalization with the ciliated cell marker FOXJ1 (pink) reveals a multicellular airway epithelium composed primarily of club cells. In *Svep1^-/-^* embryos, the bronchiolar lumina are narrowed than *Svep1^+/+^* (highlighted in white). Quantification shows an increased average of number of club cells (SCGB1A1) but not of ciliated cells (FOXJ1). (Average counts ± SEM; *t* test p < 0.05 n≤ 3). *Scale bars 200 µm or 100 µm*

The RNA-seq data from the whole RCa lobe at E18.5 revealed that genes associated with muscle development were significantly downregulated in *Svep1^-/-^* lungs, including *Acta2* (actin alpha 2) which is necessary for smooth muscle differentiation, and *Cnn1* (calponin1), a marker for mature smooth muscle (Fig. 5b). Lungs labeled with ACTA2 and particularly CNN1 confirmed the reduction in distal smooth muscle differentiation in the mutant at E18.5 (Fig. 5c) evident around the proximal airways and adjacent blood vessels, particularly CNN1 expression (Fig. 5c). The expression of ACTA2 was noticeably reduced also in lobular bronchioles in *ex vivo* organ culture of E12.5 *Svep1^-/-^* lung explants that were allowed to grow in vitro for six days (Fig. 5e) confirming that *Svep1* is necessary for proper peribronchial smooth muscle differentiation which is defective in the lungs of *Svep1* mutant embryos before saccular development.

To investigate lung airway development in *Svep1^-/-^* mutants further, we examined cellular differentiation of the proximal epithelium by staining with the proximal epithelium progenitor SRY-box 2 (SOX2) marker at the end of branching morphogenesis (E16.5)(Fig. 5f) and with both club cell secretoglobin 1a1 (SCGB1A1) and ciliated cell forkhead box protein J1 (FOXJ1) markers at E18.5 in the proximal airway epithelium (Fig. 5g). In mutants, SOX2 protein was predominantly localized in the lung core (Fig. 5f), suggesting that the airway epithelium follows a normal developmental pattern from proximal to distal at E16.5, with most proximal epithelium undergoing typical development. SCGB1A1 staining revealed that *Svep1^-/-^* deficient embryos displayed smaller distal bronchioles, accompanied by disorganized and densely packed SCGBA1+ epithelial cells, leading to an increase in SCGBA1+ cells (Fig. 5g). In contrast, the number of FOXJ1+ cells in mutants did not appear to differ from those in wild type lungs (Fig. 5g). Although *Scgba1* mRNA expression did not significantly differ between mutant and wild-type animals, likely due to normal epithelial development in the most proximal airways, the single cell RNA sequencing data indicated that the club cell cluster was enriched in *Svep1^-/-^* lungs (Suppl Fig. 3b). These finding highlights that normal airway structure relies on functional *Svep1* as the absence of *Svep1* leads to anomalies in bronchial epithelium morphology and an increased number of club cells.

### Altered FGF signaling is a contributing factor to the abnormalities in smooth muscle development observed in Svep1^-/-^ lungs

RNA-seq data from whole lungs revealed that *Fgfr2* expression was significantly increased in *Svep1^-/-^* embryos at E18.5. *Fgfr2* is expressed in the epithelium of the normal lung with decreased expression after E17.5 ^50^. In *Svep1^-/-^* embryos, however, *in situ* hybridization showed that levels of expression of *Fgfr2* transcripts persisted in the lungs of mutants, even at E18.5, compared to levels in lung of *Svep1^+/+^* embryos (Fig. 6a). Results from RT-qPCR showed a significant increase in RNA expression levels of *Fgfr2* and other members of the FGF signaling pathway, including *Fgf10*, *Fgf9*, and *Kras,* in the lungs of *Svep1* mutants at E18.5 (Fig. 6b) but not at E14.5 or E16.5 (data not shown). *Fgfr2* has two isoforms, *Fgfr2b* and *Fgfr2c*. Only *Fgfr2b* was significant at E18.5 (Fig. 6b). Furthermore, *Svep1^-/-^* mutants showed no change in the expression levels of important genes involved in lung development, such as *Tgfb1*, *Bmp*4, and *Shh at* E14.5, E16.5, and E18.5 (Data not shown).

**Figure 6.**
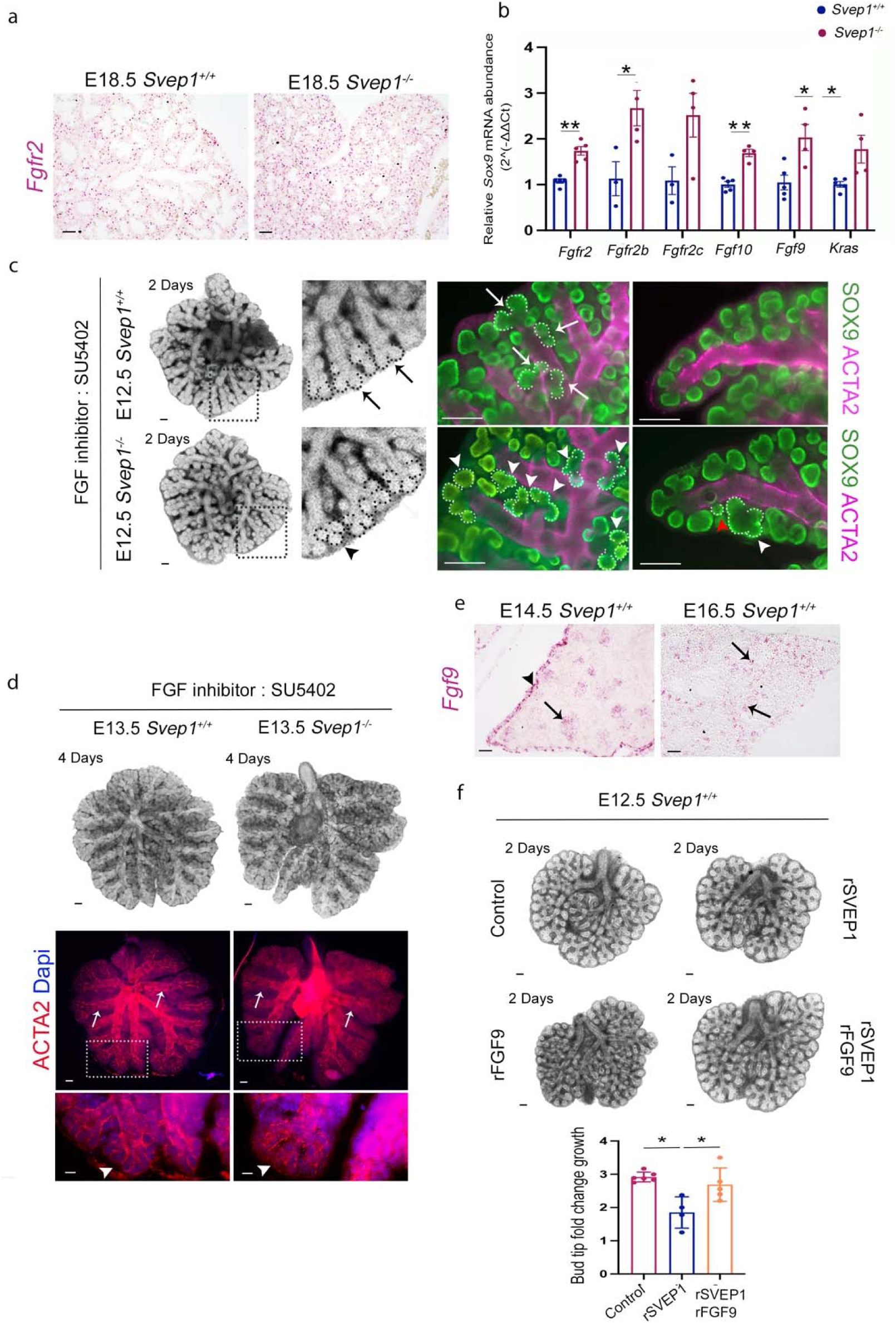
Enhanced FGF signaling in *Svep1^-/-^* embryos contributes to smooth muscle differentiation and branching defects and FGF9 rescues SVEP1 inhibition of branching. **(a)** *In situ* hybridization demonstrates increased expression of *Fgfr2* in *Svep1^-/-^* mutants. **(b)** Relative transcript abundance of *Fgfr2* (and its isoforms *Fgf2b* and *Fgfr2c*), *Fgf10*, *Fgf9,* and *Kras* showing increased expression in *Svep1^-/-^* mutant embryos at E18.5. **(c)** *Svep1^+/+^* and *Svep1^-/-^* lung explants from E12.5 embryos treated *ex vivo* with the FGF signaling inhibitor SU5402 (2.5 µg/ml). Black arrows point to normal tip morphology in *Svep1^+/+^* lungs. The black arrowhead points to distal tip trifurcations that persist in lung explants from *Svep1^-/-^* embryos. SOX9 (green) and ACTA2 (purple) staining confirm the branching anomalies (highlighted in the white dotted areas) in *Svep1^-/-^* lung explants. White arrows mark normal bifurcations, white arrowheads indicate trifurcations and distal budding; the red arrowhead highlights ectopic budding not affected by the induction of FGF. **(d)** E12.5 lung explants treated with SU5402 (2.5 µg/ml) for 4 days demonstrate reduced ACTA2 localization in distal airways of *Svep1^-/-^* lungs compared to wild type lungs. **(e)** *Fgf9* mRNA expression in wild type (E14.5) and mutant lung (E16.5). **(f)** Wild type E12.5 lung explants after 48 hours: untreated control (top left); treated with SVEP1 alone (top right); FGF9 alone (bottom left); combination of SVEP1 and FGF9 (bottom right). The combination restores a normal bud tip phenotype. (Average counts ± SEM; *t*-test p < 0.01 and <0.05, n >3). *Scale bars 100 µm (a, c, d, e)*.

FGF10/ FGFR2b signaling is necessary for airway outgrowth and promotes epithelial progenitor proliferation and migration ^52, 53^. FGF10+ cells are progenitors to parabronchial smooth muscle cells, and this progenitor status is maintained by mesothelial-derived FGF9, which inhibits smooth muscle differentiation. Smooth muscle progenitors become gradually displaced from distal to a proximal position alongside the bronchial tree in the mouse lung ^35, 54^ during development. To investigate if FGF signaling is involved in *Svep1^-/-^* branching defects and airway smooth muscle differentiation, we treated lung explants with the FGF pathway inhibitor SU5402 at a low enough dose to allow for some branching activity. SU5402 is a fibroblast growth factor receptor 1 (FGFR1) tyrosine kinase inhibitor that blocks the FGF signaling necessary for smooth muscle differentiation, leading to more smooth muscle around the airway^55^. As expected, we observed branching defects in the control and *Svep1^-/-^* lung explants after 48hr of treatment (Fig. 6c). Immunofluorescence SOX9 and ACTA2 revealed normal distal airway bifurcation in explants of *Svep1* wild type lungs and trifurcations in lung explants from mutant embryos along with ectopic branching and budding anomalies (Fig. 6c). After 4 days of FGF signaling inhibition, an increase in smooth muscle coverage around the distal airways, wrapping around the distal tips, was observed in both normal lungs and, to a lesser degree, in *Svep1^-/-^* lungs (Fig. 6d). Overall, these results suggest that the branching pattern defects in *Svep1* mutants are independent of the canonical FGF pathway, but that smooth muscle differentiation defects are partially mediated by increased FGF signaling.

### SVEP1 and FGF9 interact during branching to direct normal lung development

*Fgf9* is essential for mouse lung development, and both loss and gain of function alleles of *Fgf9* result in branching defects ^33, 35, 36, 56^. *Fgf9* is expressed in both the lung mesothelium and distal epithelium during early-to-mid branching morphogenesis. *Fgf9* levels in the mesothelium are high at E12.5, but at E14.5 *Fgf9* is limited to the distal epithelium ^33, 35^. We found that *Fgf9* is normally expressed in mesothelium and epithelium at E14.5 and exclusively in the distal airway tip epithelium of E16.5 lungs (mid-to-late branching) (Fig. 6e).

Overexpression of *Fgf9* causes branching defects that are characterized by dilatation of the distal airways ^35, 36, 53, 57^, a phenotype that is opposite to the multi-budding phenotype of *Svep1^-/-^* lungs at the pseudoglandular stage (Fig. 2c). Furthermore, overexpression of *Svep1* (Fig. 4a,b) and loss of *Fgf9* function ^33, 35^ inhibit branching. To investigate whether SVEP1 and FGF9 interact during branching morphogenesis, we treated normal lung explants with exogenous SVEP1 and FGF9 alone and in combination for two days. SVEP1 cultured explants showed a significant decrease in branching (Fig. 6f) FGF9 cultured explants exhibited dilated distal airways (Fig. 6f). Lung explants treated with both SVEP1 and FGF9 resulted in normal branching and distal airways (Fig. 6f). That the treatment with combined SVEP1 and FGF9 restored branching and distal airway morphologies suggests that an interaction between SVEP1 and FGF9 is necessary for normal airway development.

### Svep1 is necessary for cellular differentiation and saccule maturation

In *Svep1* mutants, alveolar saccules are either absent or underdeveloped, indicating that there is a disruption in the normal alveolar differentiation process. Our transcriptomics data indicate an increase in expression of AT2-related genes and a decrease in expression of AT1-related genes in *Svep1^-/-^* lungs at E18.5 (Fig. 7a). Immunolabeling with AT1 markers (PDPL and HOPX) and AT2 markers (SFTPC and ABCA3) reveals reduced PDPL and HOPX1 positive domains, expanded SFTPC expression, and intensified expression of ABCA3 in the central saccules at E18.5 (Fig. 7b). Additionally, SFTPC expression correlates with SOX9, indicating an immature AT2 cell fate phenotype in mutants (Suppl. Fig. 4a). These findings underscore that *Svep1* deficiency disrupts the balance of alveolar epithelial differentiation, favoring an AT2 cell fate. This effect is particularly pronounced in the lung periphery which retains an AT2 progenitor phenotype reminiscent of the pseudoglandular stage.

**Figure 7.**
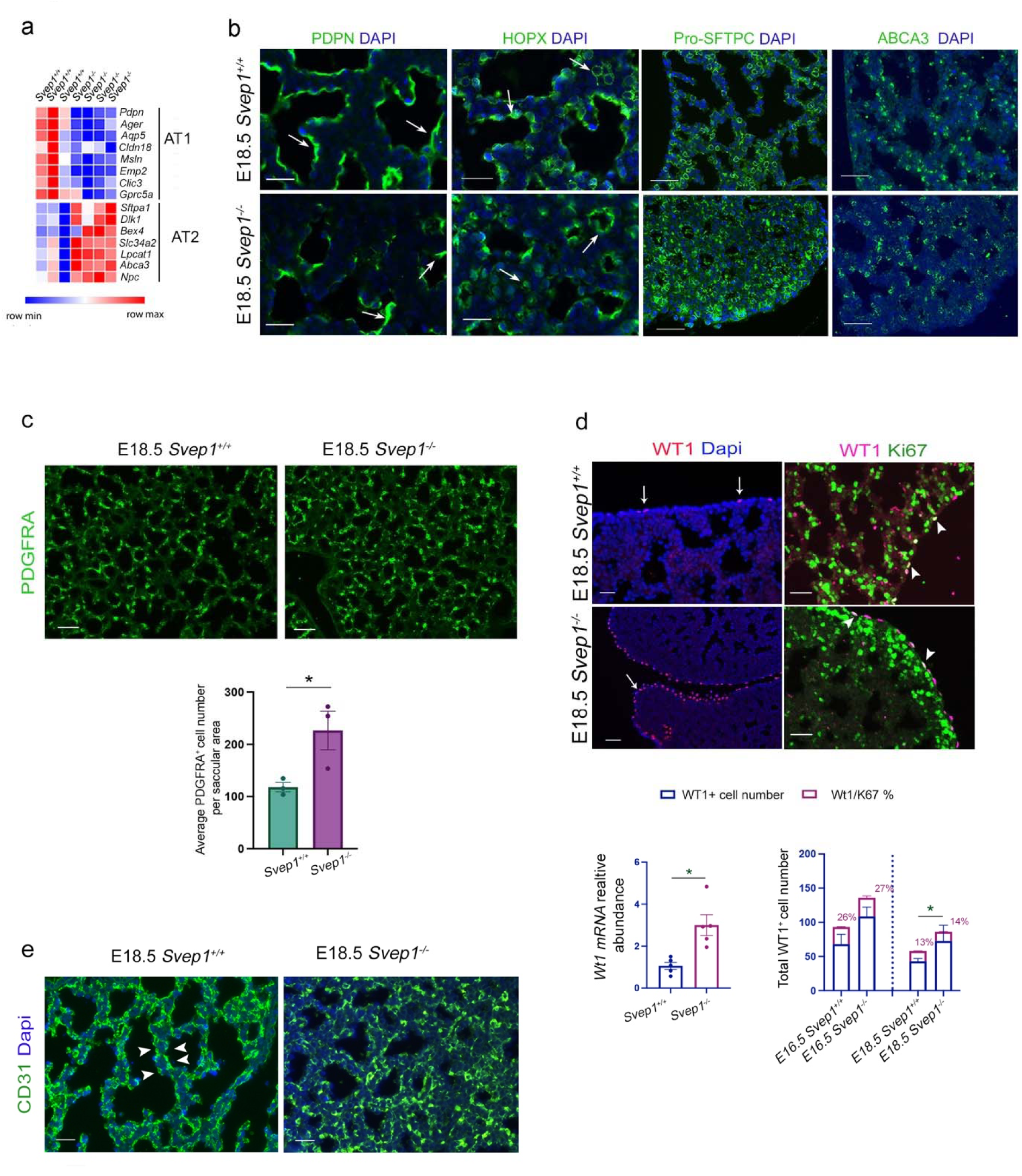
Saccular cellular and structural defects in *Svep1^-/-^* embryos at E18.5. **(a)** Expression heatmaps of selected genes associated with alveolar AT1 and AT2 cells show that *Svep1^-/-^* lungs have lower expression of AT1 genes compared to *Svep1^+/+^* lungs and higher expression of AT2 associated genes. **(b)** Immunofluorescence of PDPN, HOPX, Pro-SFPC and ABCA3 in the lungs of E18.5 *Svep 1*^+/+^ and *Svep1*^-/-^ embryos. Arrows indicate airway bud stalks. **(c)** PDGRFA+ (green) staining at E18.5. Plots of the average number of PDGFRA+ cells (average counts ± SEM; *t* test p < 0.05 n=3). **(d)** Immunofluorescence of the mesothelial progenitor marker WT1 (red) with KI67 (green) staining indicating actively dividing WT1+ progenitors (white arrows). Plot of the number of WT1+ and WT+/KI67+ cells at E16.5 and E18.5. *Svep1^-/-^* lungs have more WT1+ cells than *Svep1^+/+^* lungs at E18.5; however, the number of WT1+ KI67 lungs is unchanged between *Svep1^+/+^* and *Svep1^-/-^* lungs. (**e**) labeling of endothelial cells with CD31 illustrating the double layer of the saccular microvasculature in normal lungs (arrowhead; and disorganized microvasculature in *Svep1^-/-^* lungs at E18.5. *Scale bars 250 µm (B); 100 µm (c,d) 5 µm (e,h). Scale bars 25 µm (c,e); 50 µm (b) 100 µm (c)*.

Alveolar ACTA2 positive myofibroblasts transiently differentiate from their PDGFRA positive progenitors during septation which, during normal murine lung development, are located at the base of saccules in the regions of future alveoli, where they invaginate toward the saccule lumen ^58, 59^.To assess if *Svep1^-/-^* lungs exhibit defects in alveolar myofibroblast progenitor formation, we stained E18.5 tissues with PDGFRA, revealing a significant increase in the number of PDGFRA^+^ cells per saccular tissue in mutants (Fig. 7c). While RNA-seq analysis showed no significant difference in *Pdgfra* gene expression between mutant and wild-type lungs, the scRNA-seq data revealed a deficiency in *Pdgfra^+^* cells in the lungs of *Svep1^-/^*^-^ embryos (Suppl. Fig. 3b). The expression of *Pdgfa*, the PDGRA ligand crucial for septation, myofibroblast differentiation, proliferation, and migration ^60, 61^, was reduced in *Svep1^-/-^* embryos at E18.5 in the bulk RNA-seq data (Log2(FC)= - 0.19318, adjusted p value < 0.05 t test). PDGFRA+ cells, which are known to give rise to alveolar lipofibroblasts characterized by lipid-filled vesicles, exhibited no differences in *Plin2* (*perilipin 2*, also known as adipophilin; *Adfp*) expression between mutant and wild-type lungs at E18.5 (Suppl. Fig. 4b). These findings show that *Svep1* expression is essential for regulating alveolar myofibroblast progenitor formation. Notably, despite the absence of primary septation, *Svep1^-/-^* mutants exhibit a broad myofibroblast progenitor fate (PDGFRA+ cells). This increase may arise from an augmented number of distal airways and/or defects in the myofibroblast differentiation process itself.

The lungs are covered by a monolayer of flat mesothelial cells that differentiate from round Wilms’ tumor transcription factor 1 (WT1) positive cells that surround the lung buds during early branching ^62^. *Wt1* expression normally decreases during development ^62^ whereas the hypercellular mesothelium of the *Svep1^-/-^* lungs were enriched in WT1+ mesothelial progenitors and had increased *Wt1* gene expression (Fig. 7d and Suppl. Fig. 5). To determine if the expansion of mesothelial cells was due to an increase in cellular proliferation, we co-labeled with WT1 and KI67 (Fig. 7d). Although there was a significant increase in the number of WT1+ cells in *Svep1^-/-^* lungs (Fig. 7d, left), the percentage of WT1+ that were also KI67+ was not significantly different between normal and mutants at E16.5 (Fig. 7d, right), indicating that the disruption of mesothelial differentiation in mutants is not due to an increase in proliferation.

RNA-seq data from the whole lung revealed that vascular development genes are downregulated in the lungs of *Svep1^-/-^* embryos compared to wild-type controls (Fig. 6a). To determine, therefore, if lung microvasculature formation is affected in *Svep1^-/-^* embryos, we stained E16.5 (Suppl. Fig. 5) and E18.5 (Fig. 7e) lungs with PECAM1 (CD31). At E16.5 lung microvasculature looks normal (Suppl. Fig. 5); however, at E18.5, the microvasculature was disorganized and failed to form the double layer pattern characteristic of the saccular stage (Fig. 7e). These results indicate that *Svep1* is necessary for lung microvasculature plexus formation, which is consistent with previous findings that *Svep1* plays a role in lymphatic vascular development and remodeling in mice and explains why disruption of *Svep1* results in severe edema in embryos ^46, 47^.

## Discussion

While previous studies have led to important insights into the genetic and molecular processes that underlie lung development, our understanding of the factors that regulate the bronchial tree’s stereotypic structure and the subsequent transition from branching to saccular formation remains incomplete. The study reported here revealed, for the first time, that *Svep1* function is necessary to preserve the stereotypic pattern of the bronchial tree, contributes to the transition required to end branching morphogenesis, and promotes alveolar differentiation and maturation (Fig. 8). The results also suggest that *Svep1* regulates FGF signaling pathway for smooth muscle differentiation and that the interaction between SVEP1 and FGF9 is essential for normal airway development. Although further investigation is needed to uncover molecular mechanisms, the findings reported here provide new insights into the role the extracellular matrix plays in governing lung morphogenesis.

**Figure 8.**
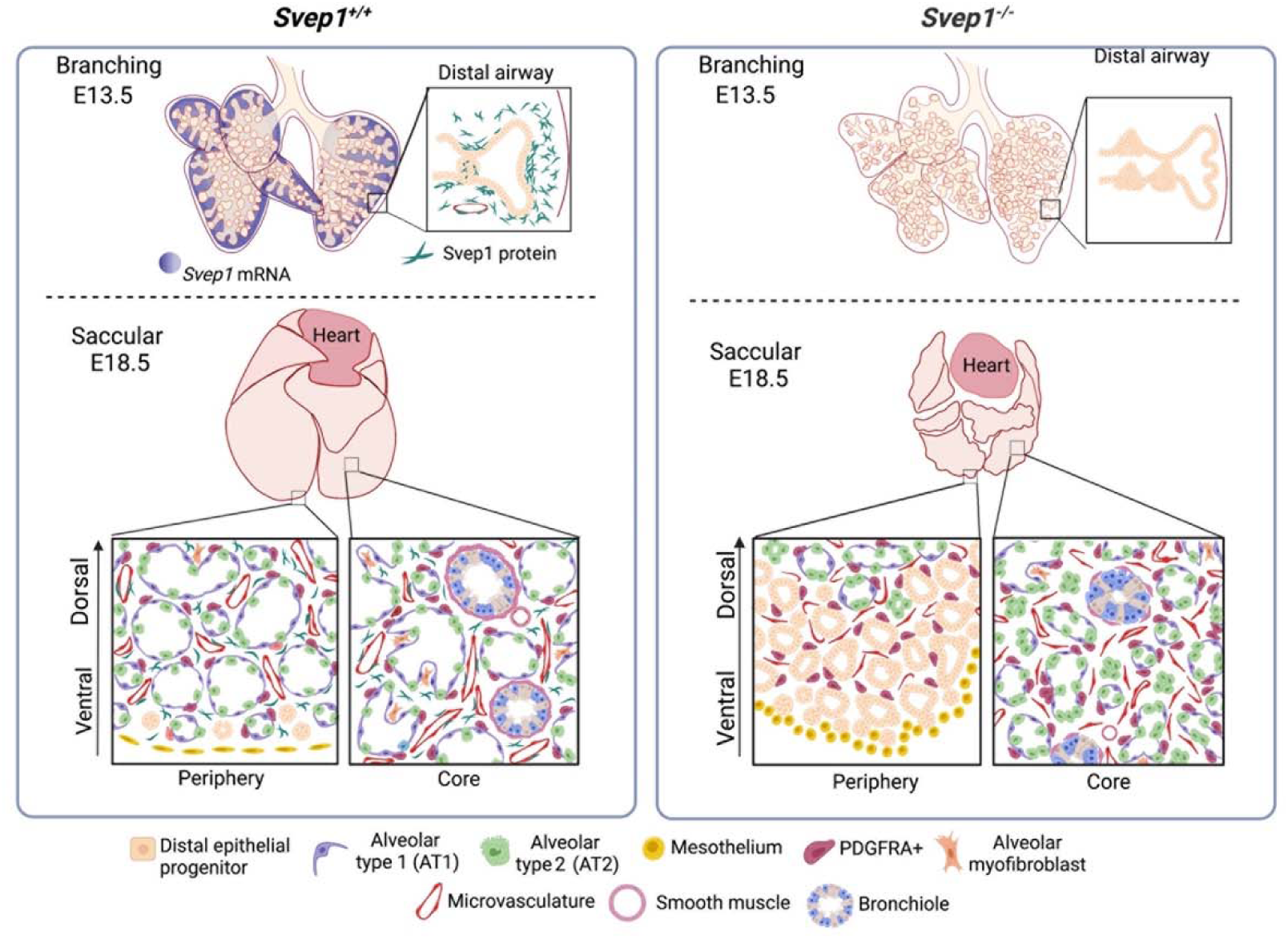
Graphical summary o normal and *Svep1^-/^*^-^ embryonic lung phenotypes during branching morphogenesis and saccular formation. (Created with BioRender©).

Our work shows that *Svep1* plays a significant role in regulating the branching program in the developing lung. The loss of *Svep1* expression leads to disordered airway tips, particularly in the distal regions where *Svep1* expression is normally high. Our results demonstrate that SVEP1 acts as a negative regulator of lung branching, inhibiting the growth of new branches in lung explants. The effects of SVEP1 on the lung bud epithelium suggests a role in shaping the lobes by restricting the potential for aberrant branching. We hypothesize that early in lung development, the local genetic program driven by FGF from its signaling centers is sufficient to direct local branching; however, as lung airway complexity increases, this local signaling becomes diffuse and may activate the entire susceptible SOX9^+^ directed epithelial branching. Therefore, negative regulators, like SVEP1 act to restrict excessive branching while ensuring proper airway patterning. Notably, at the pseudoglandular stage, lungs of *Svep1^-/-^* embryos show no significant differences in the expression of essential branching genes. This suggests that *Svep1* may not be the primary regulator of the overall genetic branching program but rather a modulator of the timing and extent of branching events. *Svep1* most likely directs branching through localized signaling without significantly impacting genes fundamental for branching morphogenesis *(e.g., Fgf10*, *Shh*, and *Bmp4*), allowing early lung developmental processes while maintaining later appropriate branching morphology.

The recovery of SVEP1 branching patterns in lung explant cultures following exogenous treatment of FGF9 suggests a potential physical interaction between SVEP1 and FGF9 in the regulation of branching similar to the SVEP1-ANG2 during the remodeling of lymphatic vessels ^47, 63^. It is plausible that SVEP1 interacts with FGF9 signaling locally in a particular distal epithelial progenitor to modify the overall branching program..

The disordered airways in the lungs of *Svep1^-/-^* embryos disrupts the shape of the lung lobes. The disrupted branching program manifests as ectopic budding and trifurcations of distal airways. The ordinarily thin and flat margins of the lung are rounded, and the tips of the lungs are foreshortened in *Svep1^-/-^* embryos. Our 3D model of the right auxiliary (RAc) lobe showed that branching in *Svep1^-/-^* embryos results in a pattern of dense rosettes generated at multiple branching sites, resulting in a cauliflower-like appearance of the lungs at E18.5. In the lungs of *Svep1^-/-^* embryos, we observed a dorsal-to-ventral maturation gradient with small immature saccules on the dorsal side and narrow airways on the lobulated ventral side. The dorsal-ventral spatial variation in aberrant lung lobe morphology is consistent with previous work showing that tissue geometry and biomechanical forces influence the genetic branching program during lung development ^7, 64^. Our data indicate that the lobe-specific phenotypes in *Svep1^-/-^* lungs are due to a complex interplay of genetic and physical factors and that the ventral lung needs tighter control than the dorsal lung to develop properly to fill the chest cavity together with the heart and diaphragm (ventral) and the body wall (dorsal). Hence, *Svep1* is especially crucial for preserving the basic structure of lung tips and edges during development.

The lungs of *Svep1^-/-^* embryos exhibit impaired cellular differentiation and saccular maturation. *Svep1^-/-^* pups die shortly after birth, with a lung morphology more typical of the pseudoglandular stage with enrichment of SOX9+ progenitor cells. The alveolar epithelium in the lungs of *Svep1* mutant embryos is biased toward AT2 cells, which correlates with the proliferative SOX9+ cell population which, in turn, is maintained by the persistent expression of genes involved in FGF signaling that impedes differentiation of AT1 cells ^27, 60^. Furthermore, *Svep1^-/-^* lungs are deficient in HOPX+ and PDPL+ AT1 cells with reduced lung bud stalks potentially leading to proximal airway hypoplasia. *Svep1^-/-^* lungs have reduced smooth muscle differentiation that can be rescued by the inhibition of FGF signaling. The observed increase in SCGBA1+ club cells and the concomitant reduction in bronchiole lumen diameter point to a profound influence of the loss of *Svep1* on airway morphology, in addition to the smooth muscle development defects around the proximal airways. Furthermore, *Svep1^-/-^* lungs show reduced expression of genes related to smooth muscle, including *Acta2*. This deficiency may contribute to the impact of the loss of *Svep1* on septa and airway hypoplasia, influencing the distalization of the airway.

Other studies have found that mechanical pressures, in conjunction with growth factors, are essential for proper alveolar differentiation during development ^65^. *Svep1^-/-^* embryos are characterized by chest cavity edema secondary to lymphatic defects at E18.5 which may cause atypical internal lung pressures that could, in turn, contribute to the observed aberrant cell differentiation. Mesothelial cells and alveolar myofibroblasts at E18.5 in *Svep1^-/-^* are also maintained in a progenitor state of immature saccular development marked by increased WT1 and PDGFRA expression in mutants.

Maintenance of the extracellular matrix has been shown previously to be essential for normal organ and epithelial tube morphogenesis ^38, 66^ and cellular differentiation processes ^66^. Our transcriptome analysis indicated that the absence of *Svep1* disrupts the expression of genes associated with cell adhesion, cell motility, and ECM organization. Our transcriptomics data showed that *Col1a1* and the basement membrane collagen *Col4a4*, are significantly reduced in E18.5 *Svep1^-/-^* lungs, disruption of which may contribute to the proliferation and differentiation defects that we observed in the lungs of *Svep1^-/-^ embryos in this study*.

The transition from branching to differentiation in lung development is essential for establishing the foundation for alveologenesis and has significant clinical ramifications for neonates born with such respiratory deficiencies. Our findings demonstrate the importance of the ECM in directing airway architecture and the transition from branching programs to alveolar differentiation. Additional research into the specific role that *Svep1* plays in preparing the lung for alveologenesis is required to understand more fully the molecular, cellular, and mechanical mechanisms critical to this transition and its potential for targeted therapeutic interventions to improve the outcomes of patients born with impaired or delayed lung development.

## Methods

### Svep1 mutant mice

*Svep1* mutant mice (B6N(Cg)-*Svep1^tm1b(EUCOMM)Hmgu^*/J; MGI:5509058) were generated by the Knockout Mouse Phenotyping Program (KOMP2) by inserting the L1L2_Bact_P cassette upstream of exon 8 of *Svep1* ^49^. This cassette contains a flippase recombinase target (FRT) site followed by a lacZ sequence and a loxP site. The first loxP site is followed by a neomycin resistance gene, a second FRT site, and a second loxP site. A third loxP site is inserted downstream of exon 8. The construct was introduced into embryonic cells and embryonic stem cell clone HEPD0747_6_B06 was injected into B6(Cg)-*Tyr*^c-2J^/J blastocysts. Resulting chimeric males were bred to C57BL/6NJ female mice to generate heterozygous tm1a (i.e., knockout first) animals^67^. No homozygous mice were recovered at weaning from heterozygous intercrosses. Heterozygous tm1a mice were then bred to B6N.Cg-Tg (*Sox2*-cre)1Amc/J mice to remove the floxed neomycin sequence and exon 8 of *Svep1*. Resulting offspring were bred to C57BL/6NJ mice to remove the *cre*-expressing transgene. Genotyping was performed as described in Supplementary Methods.

### Histology

Mouse embryos were euthanized by decapitation at E14.5, E16.5, or E18.5. Tails were collected for genotyping. Whole lungs were excised surgically and fixed in 4% paraformaldehyde (PFA) and then embedded in paraffin. Paraffin tissue blocks were sectioned (5 µm) and stained for Hematoxylin and Eosin (H&E) using standard procedures.

### In situ hybridization (ISH)

For whole mount *in situ* hybridization of *Svep1*, a 725bp segment of the *Svep1* transcript (NM_153366.4; 746-11179) was PCR amplified (PCR Master Mix, Promega). *In situ* hybridization was performed using methods published previously^68^ and developed using BM purple AP substrate (Roche) per the manufacturer’s instructions. 5 µm thick sections of paraffin embedded lung tissues were used for ISH of *Fgf10* (NM_446371, 862-1978 nt), *Fgf9* (NM_013518.4, 27-1198 nt), *Fgfr2* (NM_010207, 2-1677nt), and *Svep1* (NM_022814.2, 2879 - 3726 nt) using RNAscope 2.0 Red Detection Kit (Advanced Cell Diagnostic) according to the manufacturer’s instructions.

A *Svep1* riboprobe was synthesized with one pair of exon-exon boundary overlapping primers (5’ TGTGGTCCTCCAAGTCACGTA 3’; 5’ CCAGCAGACAGCAGAGTATGT 3’) designed using NCBI’s Primer-BLAST^69^. The purified PCR fragment was cloned into the pCR™II-TOPO® TA vector (TOPO® TA Cloning® Kit, Dual Promoter, ThermoFisher), and transformed into One Shot® TOP10 Chemically Competent Cells, ThermoFisher). Ampicillin resistance transformed colonies were selected. The orientation of the insert after linearization of the vector with SpeI restriction digestion (New England Biolabs, Inc.) was determined by Sanger sequencing. Sense and anti-sense Digoxigenin-11-UTP labeled probes (DIG RNA Labeling Mix, Sigma-Aldrich) were synthesized with SP6 and T7 RNA polymerases, respectively.

Immunohistochemistry (IHC) on sectioned lung tissue was performed using standard techniques. Details on the antibodies used for IHC are provided in Supplementary Methods. Antigen retrieval was achieved by heat treatment in a microwave oven for 20 mins at low power in 0.01 M sodium citrate buffer at pH 6. For tissues imaged using fluorescence, Alexa Fluor secondary antibodies were used (Life Technology). Images were obtained with a Nikon80i or Keyence (BZX800) microscope.

### Quantification of lung distal epithelial tips

Fixed whole lungs were washed with PBS and permeabilized in 0.5% Triton-X, 5% BSA in PBS for 1 hour at room temp. SOX9 primary antibody diluted in 1X PBS was incubated overnight at room temperature. Samples were washed and incubated with an Alexa fluor (Life Technology, USA) secondary for 2 hrs at room temperature, mounted with DAPI mounting medium. and fixed with 4% PFA in PBS for 2–4 hrs.

### Tissue-clearing for evaluation of branching patterns

Fixed whole lungs from E13.5 embryos were dehydrated in a methanol series, washed with 20% DMSO in methanol, and then re-hydrated with a methanol series. Lungs were then permeabilized with 0.2% Triton-X in PBS. Lung tissue clearing was performed using the Visikol® Histo^TM^ kit (Visikol, Inc; NJ) according to the manufacturer’s instructions. Lungs mounted in silicone-imaging chambers were imaged with a Keyence (BZX800) microscope.

### Whole mount branching immunofluorescence

Lung explants were fixed with 4%PFA, dehydrated in a PBS/methanol series, bleached with 6% hydrogen peroxide (H1009, Sigma) in methanol overnight at 4° C, then re-hydrated in a methanol series. Samples were blocked in PBS + 0.3% Triton X-100 + 5% normal goat serum for 2 hrs and incubated with primary antibodies into blocking solution overnight at 4° C. Samples were washed with PBS + 0.1% Triton X-100 + 0.1% Tween-20, followed by addition of Alexa Fluor antibodies (Life Technology) diluted in PBS + 0.3% Triton X-100 at 4 °C overnight. Samples were mounted with DAPI mounting medium. washed, then fixed with 4% PFA in PBS for 2–4 hrs. Whole lungs were imaged using a Keyence microscope. A Z-stack image series (BZ-H4XD) was generated for quantification and analyses of branching patterns of embryonic whole lungs. Time lapse videos were generated to trace the lineages of airway branches for orthogonal and planar bifurcation, and for quantification of trifurcation and ectopic branching. Statistical significance was evaluated using a two-tailed t test with significance at p <0.05.

### Embryonic lung explant imaging

Lungs from E12.5 or E13.5 embryos were dissected and transferred to the inserts of Corning transwells (Millipore Sigma) containing 2 ml of DMEM/F12 medium supplemented with 1 U/ml penicillin-streptomycin. Lungs were incubated in the chamber and time-lapse images were captured every 10 min over 48 hrs using the Keyence BZX800 software (BZ-H4CXT).

### Morphometric analysis of treated lung explants

Lung explants from E12.5 or E13.5 *Svep1^+/+^* and *Svep1^-/-^* embryos were treated for 2 days with a 146 amino acid recombinant protein antigen of *Svep1* (10 µg/ml, NP_0735725.2, region 874-971, Novus Biologicals), for 2 to 4 days with and without FGFR2 kinase inhibitor (2.5 µM SU5401, Pfizer) signaling and for two days with FGF9 (200 µM; R&D system) or FGF9+SVEP1 recombinant protein

The fold change of epithelial branches was determined by manual counting of individual branches compared to epithelial buds emerging from the lung epithelium at various intervals throughout a time lapse experiment using ImageJ ^70^. Lung buds were categorized as peripheral branches only if they appeared at the most distal end of the secondary bronchi. Quantification was performed for the left lobe only. Statistical significance of the counts was evaluated using a two-tailed *t* test with significance at p <0.05.

### RNA sequencing and analyses

Total RNA was isolated from whole embryonic lungs at E18.5 using the RNeasy Mini Kit (Qiagen, Valencia, CA, USA) according to the manufacturer’s instructions. Double stranded cDNA was synthesized from total mouse lung RNA using the iScriptTM cDNA Synthesis Kit from BIORAD following the manufacturer’s instructions. RNA samples were sent to Novogene Co. Ltd for sequencing on the Illumina platform (see Supplementary Methods). Annotation term enrichment for pathways and function for differentially expressed genes was performed using Metascape (http://metascape.org) ^71^, VisuaL Annotation Display (VLAD) ^72^ (http://proto.informatics.jax.org/prototypes/vlad/), and the Gene Ontology term enrichment tool (http://geneontology.org/docs/go-enrichment-analysis/). Heat maps of gene expression were generated using the Morpheus matrix visualization and analysis software (https://software.broadinstitute.org/morpheus).

### Single cell RNA-seq (scRNA-seq) sequencing and analysis

Single cell suspensions for scRNA-seq were prepared from E18.5 embryos, sequenced, and processed as described in Supplementary Methods. The Seurat software package was used to analyze the processed data ^73^. To determine the cell populations most impacted by the loss of *Svep1*, the distribution of cell populations in each experimental group was determined by calculating the ratio of each Seurat cluster to total number of cells from an experimental group. A t-test of the ratios was conducted to determine populations significantly enriched or depleted in the mutant tissue. Clusters demonstrating a segregation between mutant and wild type origin cells were compared to the nearest-related clusters using the Wilcoxon Rank Sum test.

### Quantitative PCR

Real-time PCR was performed on cDNA with IQ SYBR Green Supermix (Biorad,) and gene-specific primers (Supplementary Methods) and beta-actin (*Actb*) as the reference. The data were analyzed using the ΔΔCt method on the BioRad CFX manager v1.5. RT-qPCR mRNA expression was analyzed by the Wilcoxon rank-sum test.

## Data availability

RNA-seq and scRNA-seq data from this study will be submitted to a public archive upon publication. The *Svep1* mouse is available from the Mutant Mouse Resource and Research Center (MMRRC:049943-UCD).

## Acknowledgements

The authors thank Drs. Martin Ringwald and Vidhya Munnamalai for their comments on an early draft of this manuscript. Sandy Daigle (The Jackson Laboratory) provided essential technical support and advice for the scRNA-seq experiment. Pete Finger (The Jackson Laboratory) performed the toluidine blue staining.

