## Supplementary Figures for "Svep1 orchestrates distal airway patterning and alveolar differentiation in murine lung development"

**Supplementary Figure 1. Expression of *Svep1* in the developing lung of wild type C57BL/6J mouse embryos. (a-b)** *Svep1* *in situ* hybridization in E11.5 and E12.5 embryos. Arrowheads indicate high expression of *Svep1* at the distal tip of right medial (RMe) and accessory (RAc) lobes (outlined in red) and the left lobe. Arrows show *Svep1* expression along and between early airway buds (outlined in red). **(b)** Arrows mark the expression of SVEP1 (green) adjacent to epithelial cells (ECAD; purple) in E14.5 normal lung. **(c)** *In situ* hybridization showing *Svep1* expression in wild type E16.5 lungs. The area outlined in the left panel is shown at higher magnification in right-hand panel to highlight airways. *Svep1* expression is stronger at the edges of the lung (arrowheads). SVEP1 protein localized next to distal airways (arrowheads) and microvasculature (arrows). **(d)** *Svep1* protein or mRNA is seen in the mesenchyme at E18.5 and is highlighted in the primary septum. **(e)** RT-qPCR *for* *Svep1* expression showing relative fold change at E14.5, E16.5 and E18.5. **(f)** Normal and *Svep1^-/-^* embryos at E12.5, E15.5, and E18.5 show edema and cranial, limb, and tail developmental defects increasing in mutants over time. **(g)** *Svep1* mRNA is absent in E18.5 mutants. **(h)** SOX9 and KI67 mark proliferating epithelial progenitor cells in E18.5 lungs, mainly at the lung periphery in *Svep1^-/-^* mutants**.** *Scale bars: 100 µm (a,c,d,h) 200 µm (B)*

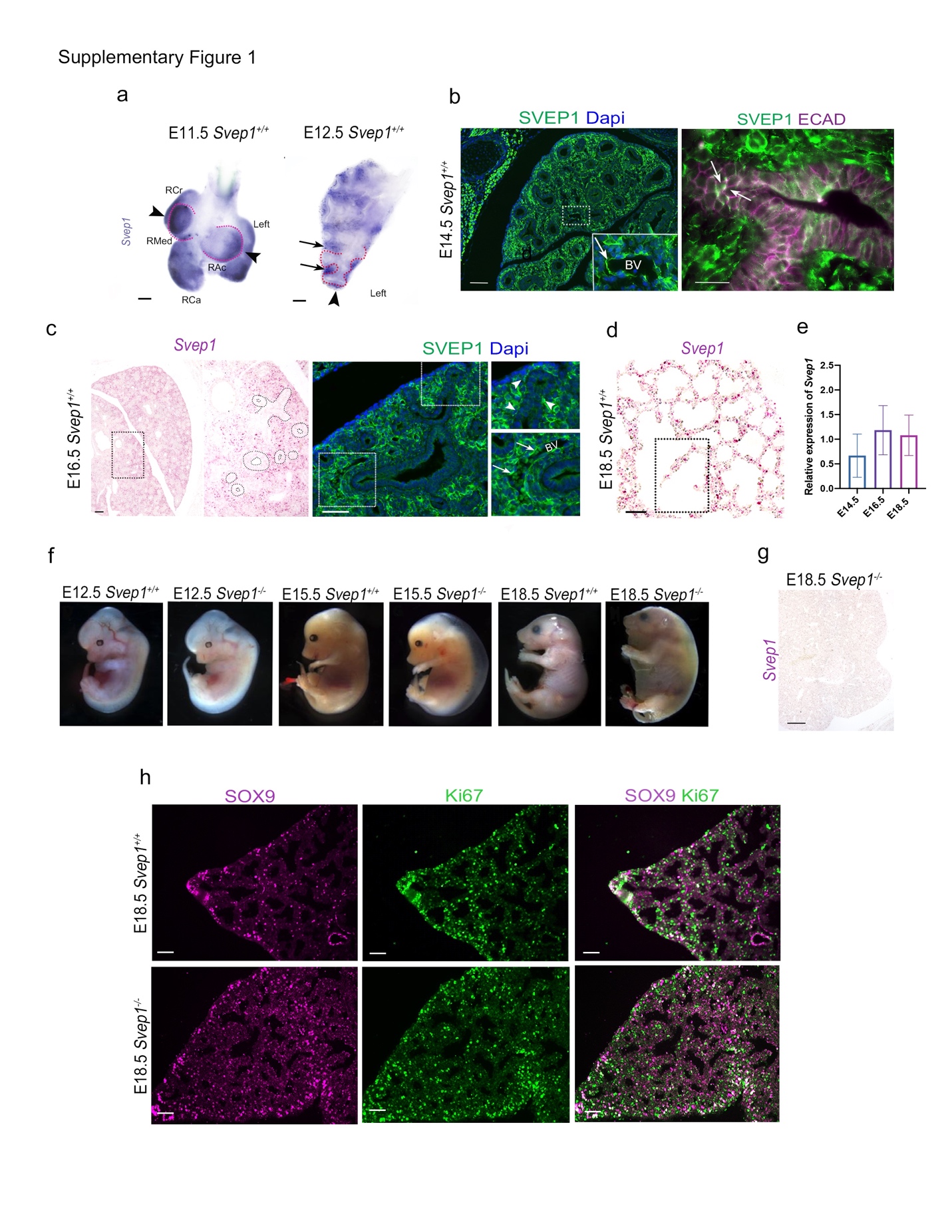

**Supplementary Figure 2. SOX9 whole mount immunohistochemistry in lung**

**lobes. (a)** SOX9 immunostaining of E13.5 lung lobe edges highlights (pink) *Svep1*

branching and bifurcation anomalies at the lung periphery. The blue highlights denote

branching domain buds. **(b)** Immunofluorescence images of E14.5 and E15.5 lung lobes showing distal airways in both wild type and *Svep1^-/-^* embryos. Arrows indicate defective branching in *Svep1^-/-^* embryos. **(c)** Graph of the ratio of length to width for

individual E14.5 lobes. *Svep1^+/+^* in pink; *Svep1^-/-^* in blue. **(d)** Graph of the ratio of length to width for individual E15.5 lobes. *Svep1^+/+^* in pink; *Svep1^-/-^* in blue. **(e)** DAPI staining of E15.5 lung lobes displays the gross morphology of the lung surface for both *Svep1^+/+^* and *Svep1^-/-^* specimens. Average lobe length-to-width ratio ± SEM; p < 0.01; n ≥ 5. Lobe abbreviations: Right Cranial (RCr), Right Caudal (RCa), Right Medial (RMe), and Right Accessory (RAc) lobes. *Scale bars:* 100 *µm*.

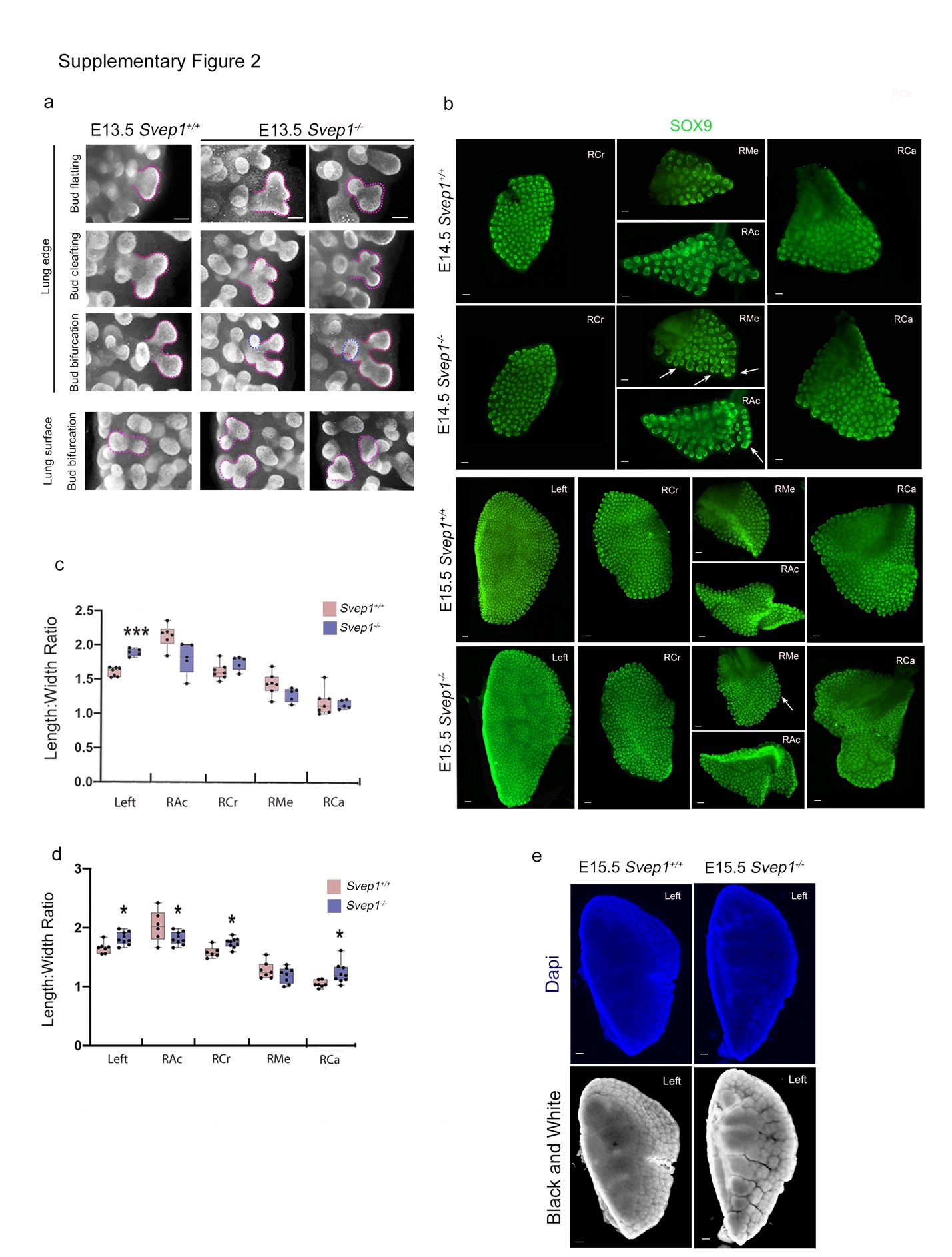

**Supplementary Figure 3. UMAP clustering of scRNA-seq generated from lungs of from *Svep1^+/+^* and *Svep1^-/-^* embryos at E18.5. (a)** UMAP clustering showing matrix fibroblast cells have the highest expression of *Svep1* (reddish orange)**. (b)** Clusters deficient in mutant cells compared to cells from wild type samples are shaded in yellow. Clusters with enrichment of cells from mutant lung samples are shaded in blue.

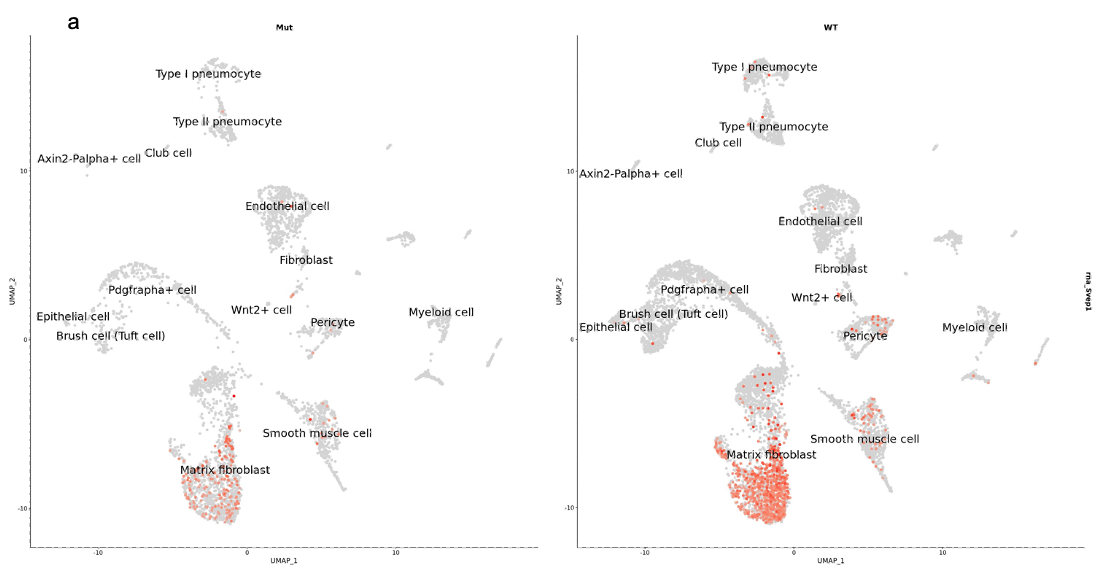

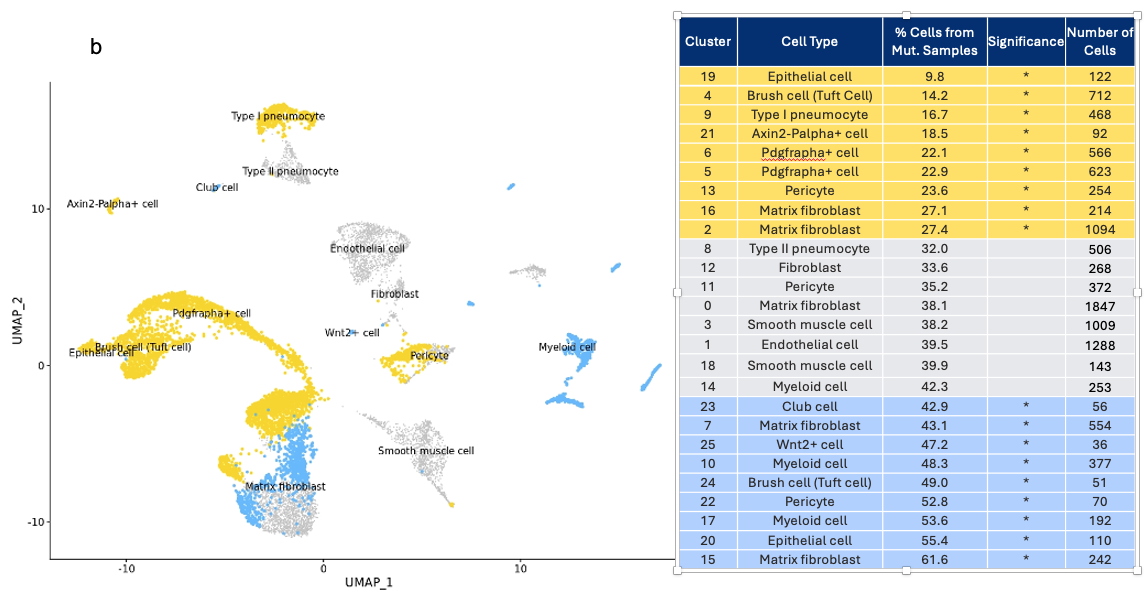

**Supplementary Figure 4. SOX9, SFTPC, and PLIN2 (ADFP) immunohistochemistry in lungs of E18.5 embryos. (a)** Epithelial localization of surfactant-associated protein C (SFTPC; green) and SRY-box 9 (SOX9; purple) expression in the distal airway tips at E18.5. Arrows indicate the colocalization of SFPC and SOX9. **(b)** Perilipin 2 (PLIN2, aka ADFP) marks lipofibroblasts in the lung mesenchyme. Expression of the *Plin2* (*Adfp*) gene in the lungs of *Svep1* mutant and wild type E18.5 embryos was not statistically different. *Scale bars:* 100 *µm*

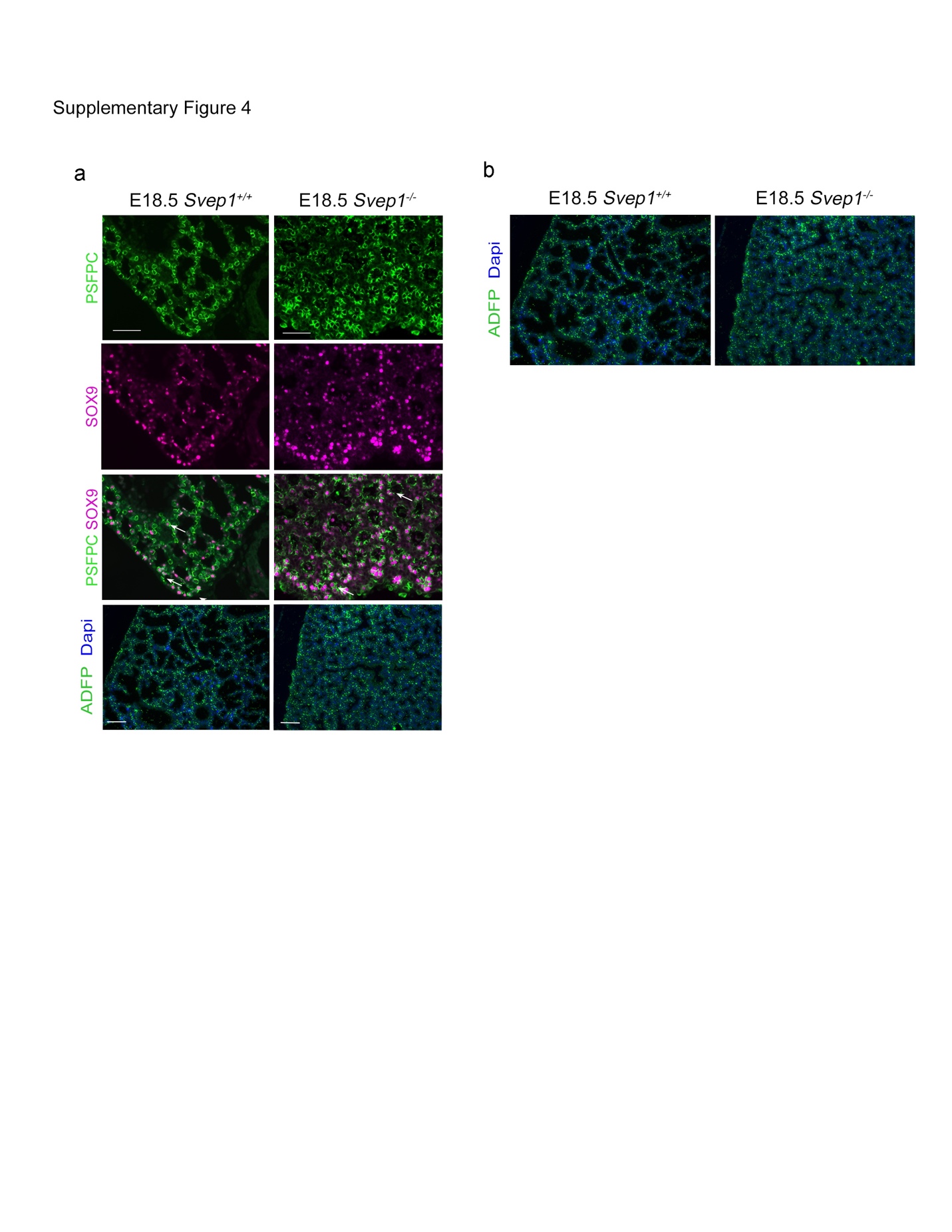

**Supplementary Figure 5. WT1 and CD31 expression in E16.5 lungs of *Svep1^+/+^* and *Svep1^-/-^* embryos. (a)** Mesothelial localization of WT1 (green) at E16.5, with arrows indicating WT1+ cells around the lung**. (b)** CD31 (green) localization marking the developing microvasculature in lungs at E16.5. Bottom panels for CD31 labeling are from regions outlined in white dotted areas in the panel above. *Scale bars:* 100 *µm*

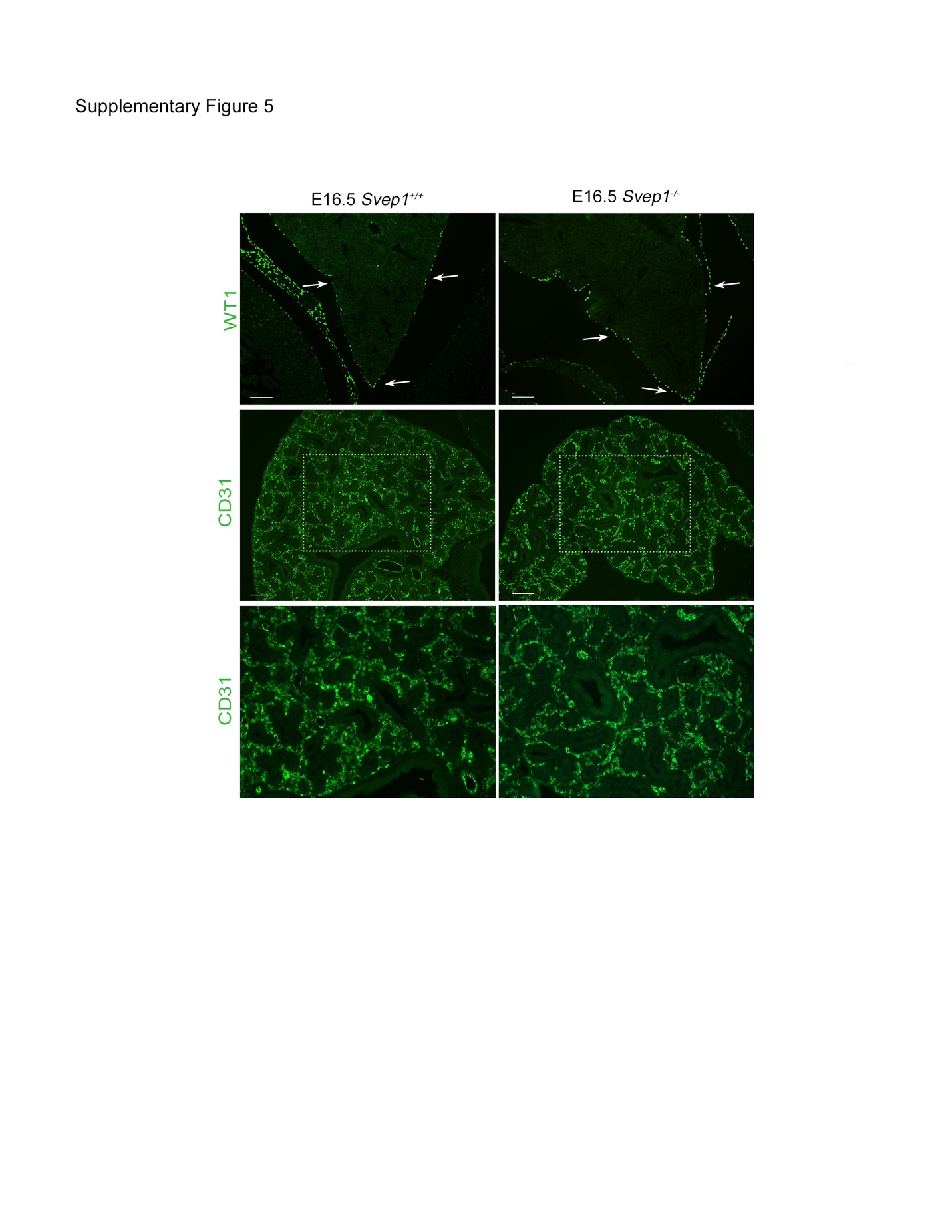
