## Supplementary Methods for "Svep1 orchestrates distal airway patterning and alveolar differentiation in murine lung development"

*Genotyping Svep1^-/-^ embryos*

Mice and embryos were genotyped by PCR using the primers shown below. The wild type (WT) PCR product is 389 base pairs (bp) in length and the mutant (MUT) product is 540 bp. DNA was isolated from tissues obtained by either ear notching or tail tipping using the Hot Shot method ^1^. Each PCR reaction consisted of 1µg of DNA, 1 µg of each genotyping primer, 4 µl of 5M betaine, 4 µl of 5X Phusion buffer, 2 µl of dNTP mix (10 mM each dNTP;) and 0.2 µl of Phusion DNA polymerase. Reactions were amplified by 25 cycles of PCR with an annealing temperature of 60^o^ C and visualized by electrophoresis on a 2% agarose gel.

| primer name | primer sequence |
| --- | --- |
| *Svep1* WT F | 5’-AGCTTTTCCACTCTAGCAAGC-3’ |
| *Svep1* WT R | 5’-CATGCAGGTCCTTTCATCCT-3’ |
| *Svep1* MUT F | 5’-CGGTCGCTACCATTACCAGT-3’ |
| *Svep1* MUT R | 5’-TCCATCCTTCAGATTTGGTCA-3’ |

*Antibodies for IHC*

The following antibodies were used for IHC: anti-α-SMA (1:200; rabbit polyclonal; Abcam), anti-α-SMA (1:200; mouse polyclonal; Santa Cruz), anti-prosurfactant protein C (1:40 0; rabbit polyclonal; Abcam), anti-prosurfactant protein C (1:50; goat polyclonal; Santa Cruz), anti-Ki67 (1:50; rat monoclonal; Invitrogen), anti-CD31 (1:50; rabbit polyclonal; Abcam), anti-PDGFR-α (1:200; rabbit polyclonal; SantaCruz), anti-podoplanin (hamster monoclonal; Abcam) , anti-SOX9 (rabbit monoclonal; Abcam), anti-E-Cadherin (1:200; mouse monoclonal; Abcam), anti-E-Cadherin (1:100; rat monoclonal; Invitrogen), anti-ERG (1:50; rabbit monoclonal; Abcam), anti-FOXJ1 (1:200; mouse monoclonal; Santa Cruz), anti-SOX2 (rat monoclonal; Invitrogen), and Uteroglobin (rat polyclonal; Abcam).

*Bulk RNA-Seq sequencing and analysis*

The cDNA library was sequenced in 2 × 150 bp paired-end layout using Illumina HiSeq2500. Sequencing data were analyzed using the Galaxy web platform (<https://usegalaxy.org/>)^2^. Sequence quality was assessed via FastQC (version 0.11.8)^3^. Sequence reads were aligned to the reference mouse genome (GRCm38) using STAR (default parameters, version 2.6.0b-1)^4^. Mapped reads were tallied using featureCounts Galaxy Version 1.6.4 (default parameters) producing a file of counted reads per gene. Differential gene expression analysis was performed using DESeq2 (version 1.22.1)^5^ in the R programming environment (version 3.5.1)^6^.

*scRNA-Seq single cell suspensions and sequencing*

To prepare the single cell suspensions, a pregnant, heterozygous female *Svep1* mouse (JAX 023814) was sacrificed at 18.5 days post coitum (dpc) by cervical dislocation (IACUC protocol 10100 to CJB). Embryos were removed from the uterine horn and transferred to dishes of warm (37^0^C) 1X Delbecco’s Phosphate Buffered Saline (DPBS; ThermoFisher, 14190250 ) and euthanized by decapitation. Tail tips from each embryo were collected for genotyping analysis. The distal tip of the right caudal lung lobe from each embryo was resected and transferred to individual wells of a 12 well plate. The methods for single-cell suspension preparation were adapted from Sekiguchi and Hauser ^7^. Each tip was submerged in a cold solution of 40 uL of 1.6 U/mL dispase II (ThermoFisher, 17105041) and 40 µL Dulbecco’s Modified Eagle Medium (DMEM)/F12 (ThermoFisher, 11039021) and incubated for 10 minutes at 37^0^C and 5% CO2. After incubation, dispase was inactivated by adding 80µL of cold DMEM/12 supplemented with 5% bovine serum albumin (BSA; Sigma-Aldrich, A8577) into each well. The lung tissues were then transferred into individual wells of a new 12 well dish, each well of which contained 80µL of calcium and magnesium free Hanks’ Balanced Salt Solution (HBSS, ThermoFisher, 14170112) to rinse off the media. Individual tissue pieces were then transferred to separate 1.5 mL microcentrifuge tubes (USA Scientific, 1415-2500), each containing 80µL protease mix (4.5µL/mL Accutase; Stem Cell Technologies, 07920; 4.5 µL/mL Accumax; Stem Cell Technologies, 07921, and 0.1mg/mL Bacillus licheniformis protease; Creative Enzymes, NATE0633, in calcium/magnesium-free DPBS). Samples were gently agitated with a pipet to dissociate for 2 minutes. Samples were then incubated on ice for 15 minutes before addition of 920µL calcium and magnesium free DPBS supplemented with 10% filtered Fetal Bovine Serum (FBS, Millipore Sigma)The samples were then filtered using Flowmi 40-um strainers (Bel-Art, H13680-0040) into individual Corning 15-mL tubes. 500µL of DPBS plus 10% FBS was added to each sample and samples were centrifuged at 112xg for 7 minutes at room temperature. Supernatants were carefully removed and cell pellets were each washed with 1mL of calcium and magnesium-free DPBS plus 1% FBS. Samples were centrifuged at 112xg for 4 minutes at room temperature then all but 40-50 µL of each supernatant was removed.

Following filtration, single cell suspensions from two mutant and two wild type *Svep1* embryos were immediately washed and resuspended in PBS containing 0.04% BSA and cells were counted in a Countess II automated cell counter (ThermoFisher). From each sample, 12,000 cells were loaded into one lane of a 10X Chromium microfluidic chip. Single cell capture, barcoding, and library preparation were performed using the 10x Chromium version 3 chemistry kit according to the manufacturer’s protocol (#CG00183). Quality checks for cDNA and libraries were conducted on an Agilent 4200 TapeStation. The cDNA and libraries were then quantified using KAPA Biosystems qPCR (Sigma-Aldrich) and sequenced using Novaseq6000 (Illumina) to an average depth of 50,000 reads per cell. A total of 6,864 cells from mutant samples and 12,309 cells from the wild type *Svep1* controls were characterized for 31,053 genome features.

Raw sequencing files were demultiplexed and FASTQ files were generated using the Cell Ranger analysis pipeline from 10x genomics (<https://support.10xgenomics.com/single-cell-gene-expression/software/pipelines/latest/what-is-cell-ranger>). Sequencing reads with mismatches within the eight-base i7 Index 1 were filtered out. The remaining libraries were aligned to the mouse reference genome assembly (GRCm38.p6) using STAR ^4^. Sequence reads with MAPQ scores less than 255 or containing bases with Q30 scores below 3 were removed. Following alignment, the cell bar codes were mapped to a list of 737,500 barcodes from 10X Genomics, with one allowed mismatch. Cell barcodes associated with a unique molecule identifier (UMI) count lower than the threshold were filtered out. A count matrix of the remaining cells and their associated genes was generated by CellRanger (10X Genomics) and used for downstream analysis. Differential gene expression was defined as genes with a p-value < 0.05 expressed in at least 25% of the analyzed populations, and -0.25 > log fold change > 0.25.

*Quantitative PCR primers*

The following primer pairs were used for qPCR of mouse genes with *Actb* as the reference for normalization. Primer information:

*Sox9*: 5’ AGGAAGTCGGTGAAGAACGG3’, 5’GGACCCTGAGATTGCCCAGA3’

*Fgfr2*: 5’ TGTGCAGATGGGATTACCGT3’, 5’ ATTTGGTTGGTGGCTCTTCTG3’

*Kras:* 5’GATGTGCCTATGGTCCTGGTA3’, 5’GCATCGTCAACACCCTGTCT3’

*Fgfr2b*: 5′AGCTCCAATGCAGAAGTGC TGG 3′, 5′ GTTTGGGCAGGACAGTGAGCC 3′;

*Fgfr2c*: 5′ CCACGGACAAAGAGATTGAGG 3′, 5′ TGTCAACCATGCA GAGTGAAAG 3′;

*Actb*: 5′ CGGCCAGGTCATCACTATTGGCAAC3′, 5′ GCCACAGGATTCCATACCCAAGAAG3′.
