## Supplementary material for "Svep1 orchestrates distal airway patterning and alveolar differentiation in murine lung development": Video Legends

Video 1

Description: E12.5 *Svep1^+/+^* lung explant culture time-lapse images over 48 hours showing normal branching morphogenesis.

Video 2

Description: Time-lapse images of E12.5 *Svep1^-/-^* lung explant culture over 48 hours demonstrating aberrant branching at the tips and edges of the lung, primarily in the form of trifurcations.

Video 3

Description: Time-lapse images of E12.5 *Svep1^-/-^* lung explant culture over 48 hours show ectopic branching on the left lobe.

Video 4

Description: Control E12.5 *Svep1^+/+^* time-lapse images for the rSVEP1 treated lungs (Video 5) showing normal branching.

Video 5

Description: Time-lapse images of E12.5 *Svep1^+/+^* lung explants treated with rSVEP1 protein for 48 hours demonstrating decreased secondary branching.
